# Living with the invader: two native deer, two outcomes along a gradient of axis deer invasion

**DOI:** 10.64898/2026.09.21.753347

**Authors:** Alexandra Cravino Mol, Santiago Mirazo, Alejandro Brazeiro, Juan Andrés Martínez-Lanfranco

## Abstract

Exotic species can reshape native communities, in the extreme, to local extirpation. Before that, natives may adjust where and when they use the landscape, in ways that allow coexistence and may be the earliest sign of trouble. Distinguishing coexistence built on such adjustment from one built on pre-existing niche differences requires comparing a native where the invader is absent with the same native where it is present, across sites at different invasion stages. Using eight years (2018–2026) of camera-trap data from 103 stations across Uruguay (154,590 camera-nights), we asked how the invasive axis deer (*Axis axis*) affects the space and time use of two native cervids: the forest-dwelling brown brocket (*Subulo gouazoubira*) and the grassland-restricted pampas deer (*Ozotoceros bezoarticus*). Activity-overlap, directional-avoidance, hierarchical activity, and two-species occupancy analyses showed sharply divergent dyads. *A. axis* and *S. gouazoubira* shared forest yet segregated in space and along the understory stratum the brocket requires and the invader avoids, and shifted their diel activity by context; understory cover fell with *A. axis* abundance, consistent with habitat modification. For *A. axis* and *O. bezoarticus*, segregation was intrinsic – opposite diel niches and habitats – with no support for a detectable shift in pampas deer behavior. The only behavioral signal was asymmetric: the invader, rather than the native, appeared to adjust, though few shared sites warrant caution. The same invader thus drives different mechanisms, but coexistence is provisional: *A. axis* is still expanding, far from equilibrium, so these shifts read best as early warning signals.

**Lay Summary:** Conservation usually waits for numbers to drop. But animals change their behavior first. Over eight years in Uruguay, we watched an invasive deer spread past two native deer: one shifted its hours and its haunts, then went undetected at one continuously monitored site; the other, active by day on different ground, showed no detectable behavioral response. Watching behavior, not just counting animals, can reveal trouble early.

## Introduction

Biological invasions are among the leading drivers of biodiversity loss worldwide, and their ecological and socio-economic costs continue to rise as global trade accelerates the rate of species introductions (Mack et al. 2000; Seebens et al. 2017; Diagne et al. 2021; IPBES 2023). Beyond outright displacement, introduced species can reshape native communities through competition, predation, disease transmission, and structural habitat modification, altering the selective pressures that resident species experience (Sakai et al. 2001; Simberloff et al. 2013; Haubrock et al. 2026). Native fauna respond to these pressures along a continuum between two extremes: coexistence with no detectable effect on the native at one end, and local extirpation at the other. Between them, coexistence can be achieved through fine-grained partitioning of shared space and time, depending on interaction strength, the behavioral plasticity of native species, and the structural attributes of the landscape (Schoener 1974; Kronfeld-Schor and Dayan 2003).

The mechanisms by which native species persist alongside an invader fall broadly into two non-mutually-exclusive categories. The first is active behavioral adjustment, in which natives modify their activity, habitat use, or movement in response to the invader, partitioning the niche along temporal or spatial axes (Frey et al. 2017; Niedballa et al. 2019). Although these behavioral changes prevent direct short-term mortality, they generate sustained physiological and demographic costs (e.g., reduced foraging efficiency, heat stress, reduced reproductive success). These behavioral modifications may function as early warning signals that precede population collapse (Wong and Candolin 2015; Cerini et al. 2023). The second is intrinsic niche segregation, in which pre-existing differences in habitat preference, activity rhythm, or trophic ecology limit overlap independently of the invader’s behavior (MacArthur and Levins 1967; Chesson 2000). These mechanisms imply contrasting futures: behavioral accommodation may erode as invader pressure intensifies, whereas intrinsic segregation predicts more stable coexistence.

Exotic ungulates are among the most frequently introduced vertebrates worldwide and a growing threat to native biodiversity (Sakai et al. 2001; Luque et al. 2014). Among them, the chital or axis deer (*Axis axis*; Aa hereafter), native to the Indian subcontinent, has been introduced for hunting across several continents and has become established and overabundant in parts of South America. In Uruguay, the species was introduced to the western littoral in 1927 – almost exactly a century ago – and has since expanded through a combination of spread from long-established western populations and secondary foci within Uruguay associated with human-mediated translocations, generating a heterogeneous invasion landscape rather than a single continuous wavefront (González and Martínez-Lanfranco 2010; Pereira-Garbero et al. 2013; Cravino et al. 2021; Cravino and Brazeiro 2023). At a regional scale, *A. axis* now thrives along the Paraná and Uruguay river watersheds of Argentina, Uruguay and southern Brazil, where it has become overabundant and difficult to control despite sustained management efforts (Pereira-Garbero et al. 2013; Gürtler et al. 2018; Burgueño et al. 2021; Gürtler et al. 2024). It was first recorded in Brazil in 2009, in the Pampa of westernmost Rio Grande do Sul (Sponchiado et al. 2011), and bioclimatic niche models predict substantial scope for further spread across the region (Etges et al. 2023). Throughout its introduced South American range, *A. axis* has been reported to compete with native deer, to modify vegetation through herbivory, and to threaten rare or declining species through habitat modification (Pereira-Garbero et al. 2013; Gürtler et al. 2018; Cravino et al. 2021; Szpilbarg et al. 2025).

This expansion overlaps the ranges of two native cervids of contrasting ecology in Uruguay. The brown brocket deer, also known as the gray or brown brocket (*Subulo gouazoubira;* Sg; formerly *Mazama gouazoubira*; Bernegossi et al. 2022) is a small, solitary, forest-dwelling species widely distributed across the Neotropics, strongly associated with woody vegetation and dense understory (Rivero et al. 2005; Andrade-Núñez and Mitchell Aide 2010; Black-Décima et al. 2010; Albanesi et al. 2019; Grotta-Neto et al. 2019; Weiler et al. 2020; Martínez-Polanco 2026). The pampas deer (*Ozotoceros bezoarticus*; Ob) is a grassland specialist whose Uruguayan populations correspond to two subspecies: *O. b. arerunguaensis*, restricted to the basaltic region of north-central Uruguay (departments of Salto, Paysandú, and western Tacuarembó) and *O. b. uruguayensis* restricted to a smaller region in the east (department of Rocha) (González et al. 2002; Cosse et al. 2009; González and Martínez-Lanfranco 2010; Cosse and González 2013; Gonzalez et al. 2023). Both subspecies are of high conservation concern. *S. gouazoubira* and *O. bezoarticus* differ fundamentally in habitat use, body size, diel activity, and degree of habitat plasticity, providing a natural contrast for testing whether they respond to the same invader through similar or distinct mechanisms.

The temporal axis is a particularly important dimension of coexistence among sympatric ungulates, which frequently mitigate competition by differing in diel activity; temporal niche partitioning is a prevalent mechanism enabling coexistence among sympatric species (Schoener 1974; Kronfeld-Schor and Dayan 2003; Kronfeld-Schor et al. 2013; Frey et al. 2017). Camera trapping has become a powerful tool for quantifying these patterns and the spatial associations that accompany them (Burton et al. 2015; Caravaggi et al. 2017; Diete et al. 2017; Niedballa et al. 2019; Cravino and Brazeiro 2023), since sympatric species must partition time or space to coexist and spatiotemporal partitioning reduces competition and the potential for agonistic encounters (Vanak et al. 2013).

Here we use eight years (2018–2026) of camera-trap data from 103 stations across Uruguay (Supplementary Material Table S2; species distributions in Supplementary Material Figure S1) to quantify how the exotic deer *A. axis* affects the spatial and temporal patterns of habitat use of two native cervids (*O. bezoarticus* and *S. gouazoubira*). Because the expansion of *A. axis* is still underway and has not reached equilibrium in distribution or abundance, we treat the coexistence we observe as a stage in an unfolding process rather than as an outcome, and we ask what sustains it. Two possibilities carry different implications. Coexistence may rest on behavioral accommodation, with natives adjusting their use of space and time in the presence of the invader; such adjustments carry physiological and demographic costs and could signal early conservation problems not yet visible in distribution or abundance. Alternatively, it may rest on pre-existing differences in habitat and activity that limit overlap regardless of the invader’s behavior, a configuration expected to be more stable. The two natives differ enough in ecology that they need not follow the same route, and the ecology of *A. axis* in Uruguay is only now being described, so we treat these as alternatives to be discriminated rather than as predictions to be confirmed. We therefore asked whether diel activity overlap between species differs across invasion levels; whether the diel activity curve of each species shifts in the presence of the other within shared sites; whether spatiotemporal avoidance is directional and consistent with return-time dynamics; what local- and landscape-scale factors predict the occurrence of each species; and whether jointly modeled occurrence reveals positive or negative spatial associations between species.

## Methods

### Study area

The study was conducted across Uruguay (33° S, 56° W), in southeastern South America. The landscape is dominated by the natural and semi-natural grasslands of the Río de la Plata region (Soriano 1991; Paruelo and Jobbágy 2007; Baldi and Paruelo 2008; Baeza et al. 2022; Gallego et al. 2024), interspersed with riparian and hill forests, wooded savannas, and increasingly extensive commercial *Eucalyptus* and *Pinus* plantations (Jobbágy et al. 2006; Paruelo et al. 2006; Gautreau 2014; Cravino and Brazeiro 2021; Baeza et al. 2022; Gallego et al. 2024). The north-central region (departments of Salto, Paysandú, and western Tacuarembó) has shallow basalt-derived soils that support extensive natural grasslands of high conservation value (Brazeiro 2015; Baeza et al. 2022; Gallego et al. 2024). The climate is humid temperate, classified as Cfa (temperate without dry season, hot summer) under the Köppen–Geiger system (Köppen and Geiger 1926; Beck et al. 2018), with rainfall distributed evenly through the year (∼1,200 mm annually) (INUMET 2020).

In Uruguay, 12 study sites were monitored, and each was assigned an invasion level describing how far the national expansion of *A. axis* has progressed locally. Assignment was based on the documented distribution and expansion history of the species in Uruguay (Cravino et al. 2021), which draws on records well beyond our camera-trap network; the relative abundance index recorded at our own stations (RAI; detections per 100 camera-nights) is reported here as independent corroboration rather than as the criterion, and it recovers the same ordering: no invasion (*A. axis* absent), low invasion (RAI = 2.4), moderate invasion (RAI = 4.0), high invasion (RAI = 10.5–13.6) and total invasion (RAI = 28.2), the last comprising sites where the native brown brocket (Sg) went undetected. Because the level describes the invader rather than the pair of species compared, it applies to both dyads: the northern sites where axis deer co-occurs with the pampas deer (*O. bezoarticus arerunguaensis* – Ob –) fall within the high-invasion level. Where Aa and Sg co-occurred, the invader accounted for 14% (low), 42% (moderate), and 81% (high) of paired detections. RAI is an imperfect proxy for density and can be influenced by among-habitat differences in detectability, so these assignments are approximate. Thus, invasion level should be read as a site-context classification that combines invasion history, local abundance and geography, not as a strictly chronological sequence replicated independently across space. No sampling sites fell within the range of *O. bezoarticus uruguayensis*. Because these contexts are defined as geographic zones, they are inherently associated with locality and, to some extent, sampling period; we therefore interpret context differences as reflecting invasion stage in this landscape rather than as effects isolated from geography, and we return to this in the Discussion. Throughout the figures and tables, each panel is identified by its invasion level and by a combination of species silhouettes indicating the dyad compared (Figures 1 and 4), and these levels framed all temporal-overlap and spatiotemporal-avoidance comparisons.

**Figure 1.**
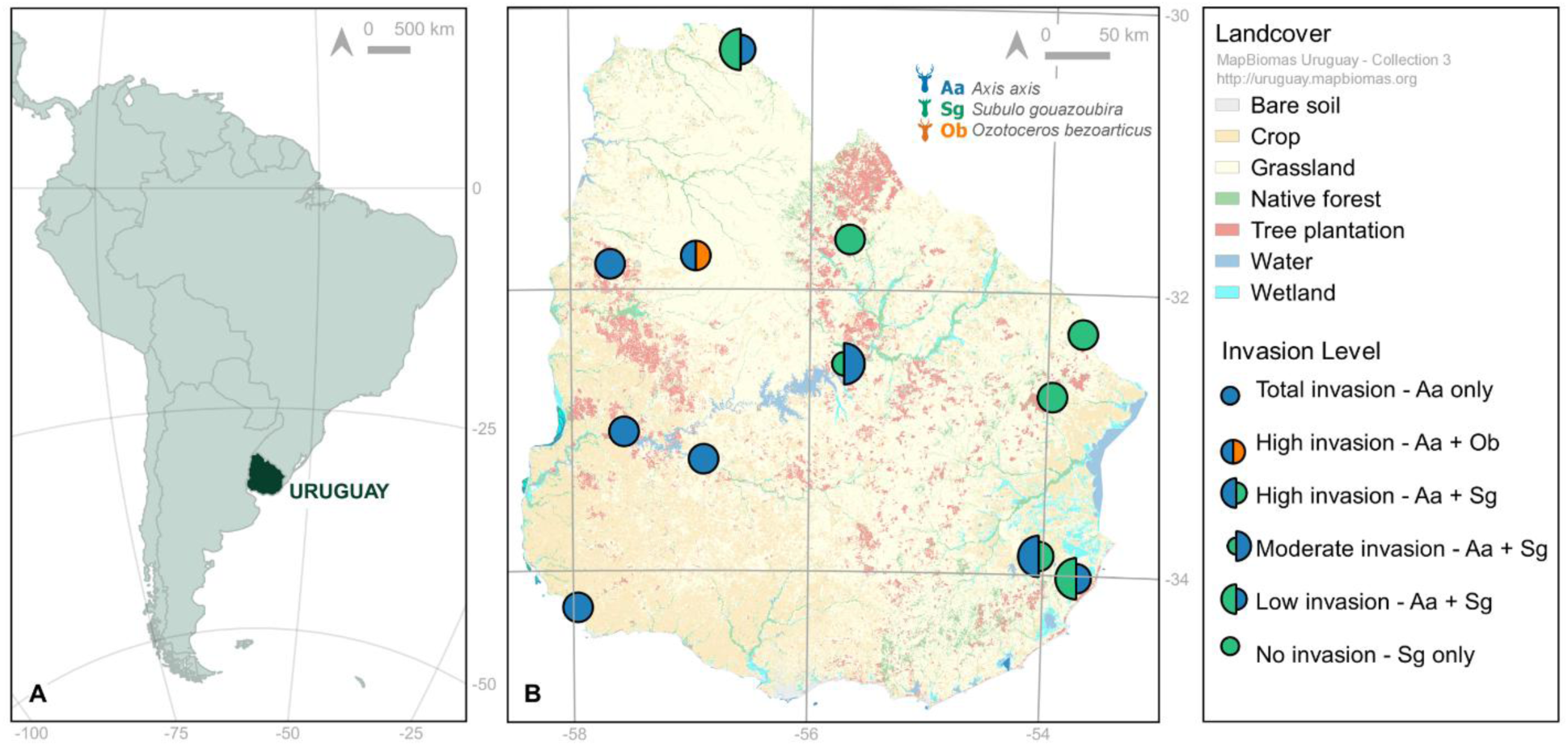
A. Uruguay within South America; B. Study areas across Uruguay. The map shows the centroid of the camera-trap stations within each area, with each site identified by the invasion level assigned to it: allopatric sites, sites numerically dominated by the invasive axis deer (Aa) -total invasion- or by the native brown brocket (Sg) -no invasion-, sites at the moderate-invasion level where the brown brocket is declining, and the high-invasion sites where the axis deer co-occurs with the pampas deer (Ob) or with Sg. Species are identified by silhouette and by color (Aa, blue; Sg, green; Ob, orange). Alt text: Two-panel figure. Panel A locates Uruguay within South America. Panel B is a land-cover map of Uruguay, shaded for grassland, crop, native forest, tree plantation, water and wetland, on which twelve circles mark the study localities. Each circle is colored by the species present and by invasion level: solid green where only *S. gouazoubira* occurs, half green and half blue at low to high invasion where both deer occur, solid blue where *A. axis* occurs alone, and half blue and half orange at the locality where *A. axis* co-occurs with *O. bezoarticus*. Invaded localities lie mainly in the west and south, uninvaded ones in the north and east.

**Figure 2.**
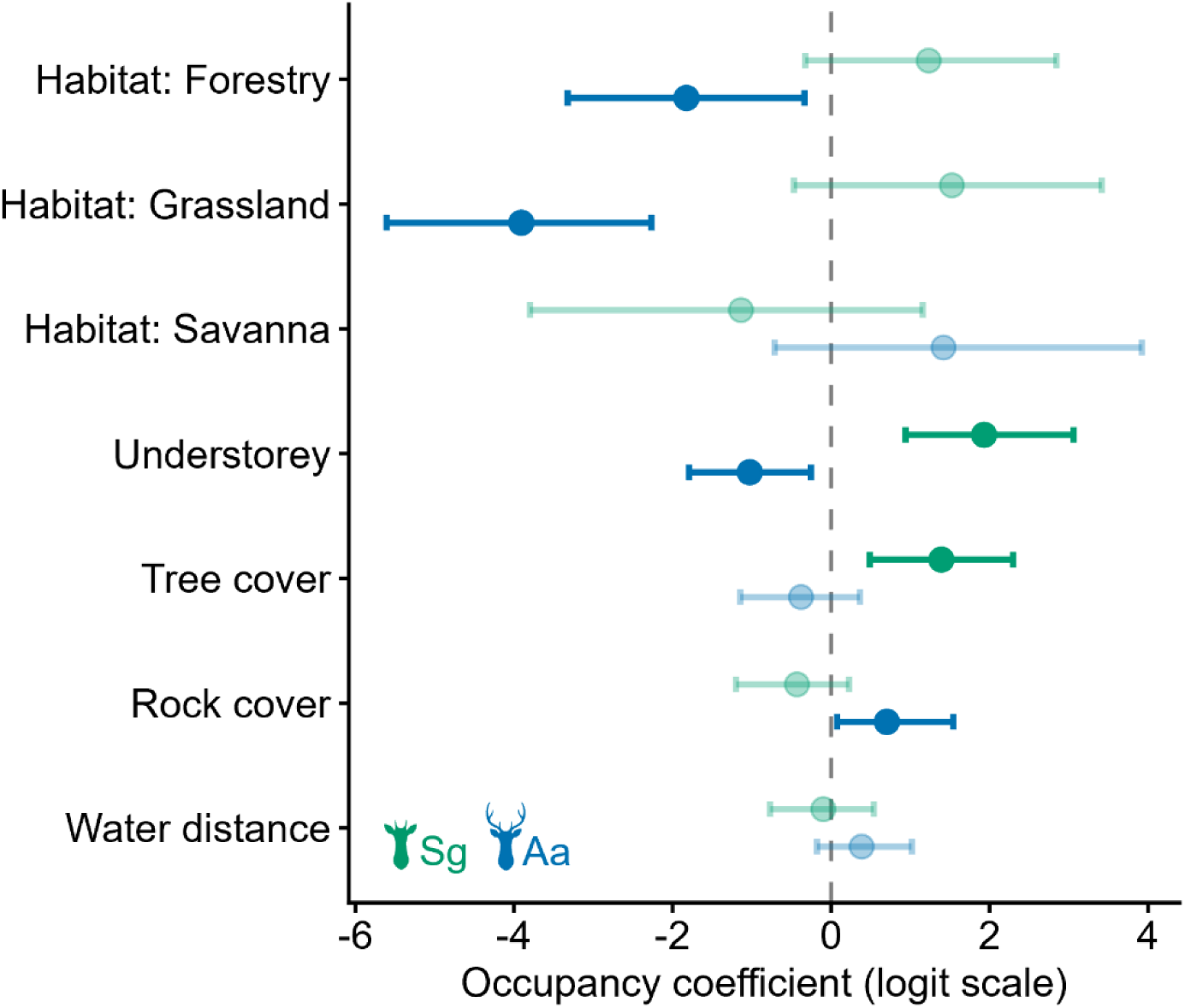
Forest plot of single-species occupancy coefficients (posterior median ± 95% credible interval, logit scale) for *A. axis* and *S. gouazoubira* (shared local-habitat covariates). Coefficients whose 95% interval crosses zero are shown faded; the dashed line marks no significant effect. The opposing understory responses of *A. axi*s and *S. gouazoubira* − negative and positive, respectively − are the clearest contrast between the two species. Species are identified by silhouette and color as in Figure 1. Alt text: Forest plot of occupancy coefficients on the logit scale for *A. axis* and *S. gouazoubira*. Seven covariates are stacked vertically; each has a point for the posterior median and a horizontal bar for the 95% credible interval, read against a dashed vertical line at zero. Faded symbols mark intervals that cross zero. The two species take opposite signs on understory and tree cover, negative for *A. axis* and positive for *S. gouazoubira*, and *A. axis* is strongly negative for grassland and forestry habitat.

**Figure 3.**
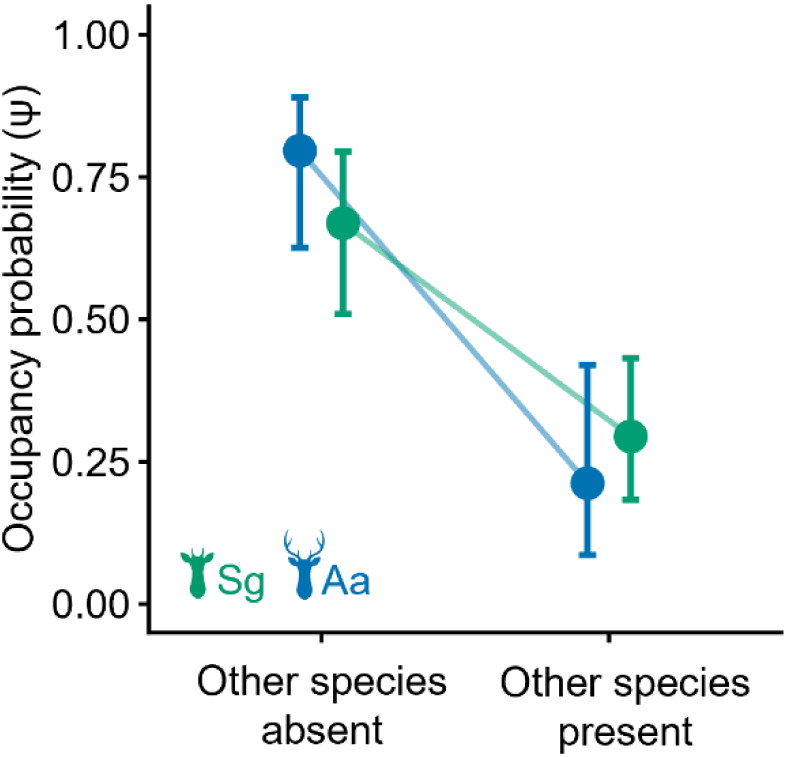
Spatial exclusion between *A. axis* and *S. gouazoubira* from the two-species co-occurrence model (occuMulti; Rota et al. 2016). Points show the conditional occupancy (ψ) of each species when the other is absent vs present, with 95% confidence intervals (bootstrap). Both species show a marked decline in occupancy in the presence of the other (*A. axis* from 0.80 to 0.21; *S. gouazoubira* from 0.67 to 0.29), reflecting the negative interaction term (f3 = -3.69, p < 0.0001) that persists after controlling for continuous habitat covariates. Species are identified by silhouette and color as in Figure 1. Alt text: Line-and-point plot of conditional occupancy probability. For each species, two points joined by a line compare sites where the other species is absent with sites where it is present, with vertical 95% confidence bars. Both lines fall steeply: *A. axis* from about 0.80 to 0.21 and *S. gouazoubira* from about 0.67 to 0.29.

**Figure 4.**
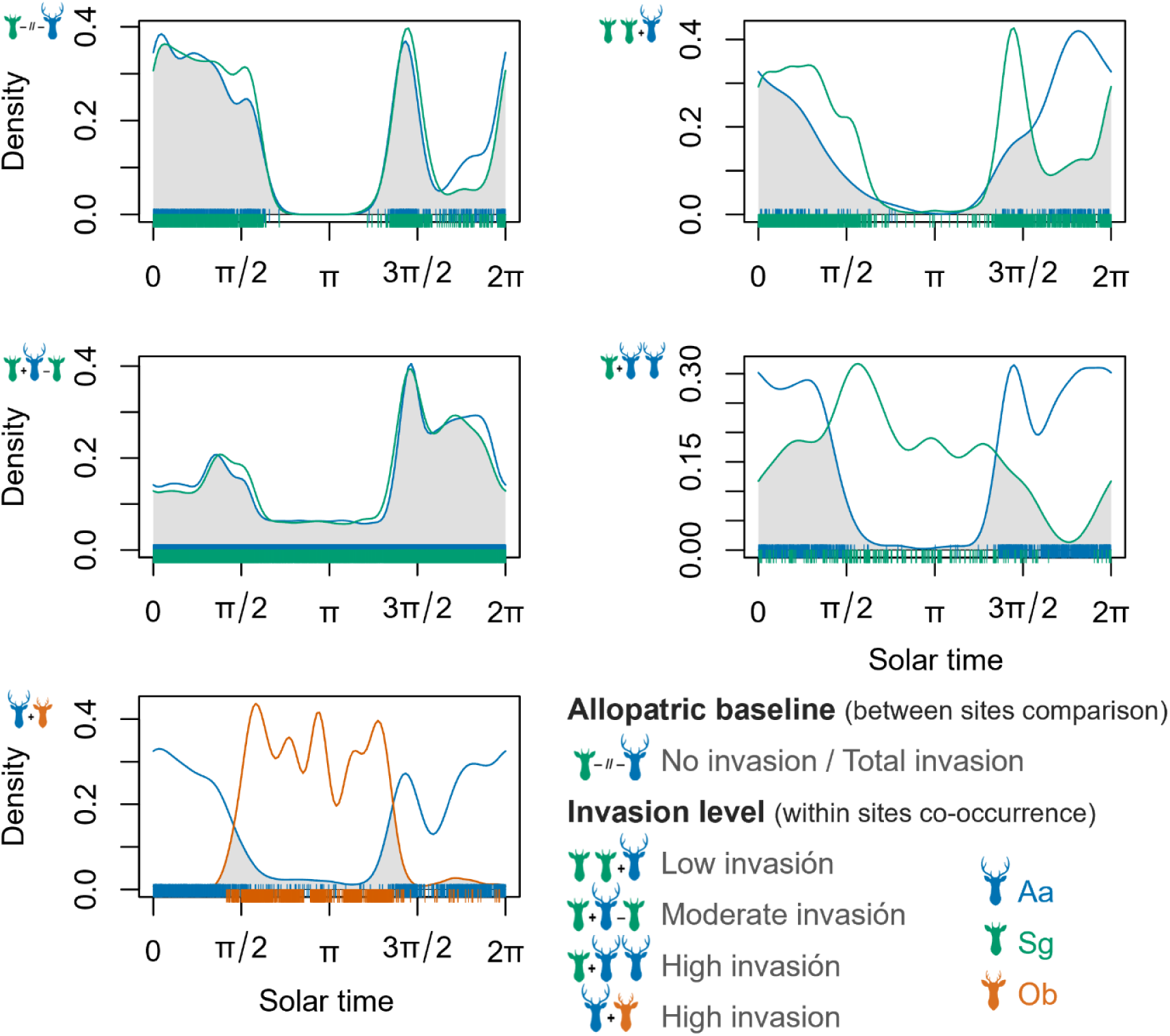
Diel activity patterns of *Axis axis, Subulo gouazoubira* and *Ozotoceros bezoarticus* across invasion levels, expressed in solar time (0–2π; sunrise ≈ π/2, sunset ≈ 3π/2). Curves are kernel density estimates of activity fitted to the solar-time distribution of detections; the shaded overlap between two curves represents the coefficient of overlap (Δ). Panels contrast each invasion level (allopatry, low, moderate, and high invasion, and the Aa/Ob dyad), with silhouettes indicating the dyad compared in each panel. Note the high overlap between *A. axis* and *S. gouazoubira* when measured in single-species contexts (Δ = 0.955), its reduction where the species co-occur, and the near-absence of overlap between *A. axis* and *O. bezoarticus* (Δ = 0.178). Species are identified by silhouette and color as in Figure 1. Alt text: Five panels of overlaid kernel density curves of activity against solar time, one panel per co-occurrence context, with the area of overlap shaded and rug marks showing individual detections. Blue curves are *A. axis*, green *S. gouazoubira* and orange *O. bezoarticus*. The two deer curves are almost coincident where each occurs alone and become progressively displaced as invasion advances; *A. axis* and *O. bezoarticus* barely overlap at all, the former being active mostly at night and the latter almost entirely by day.

### Camera-trap survey

Between January 2018 and January 2026, we operated a network of 103 motion-triggered camera-trap stations across 12 localities in 11 departments of Uruguay, deliberately sited to span the invasion gradient – from long-established, high-density *A. axis* populations in the western littoral to recently colonized or as-yet-uninvaded areas elsewhere (Figure 1). Stations followed a stratified-random design (stratified by habitat type) with a minimum separation of 500 m between stations within a stratum. Individual stations were active for 363–2,919 days (mean 1,501), accumulating 154,590 camera-nights over the study. Cameras were mounted on trees or wooden posts 30–50 cm above the ground and set to take three photographs per trigger followed by a 10-s delay, without bait or lures. Each station was characterized by 11 local- and landscape-scale environmental variables: dominant habitat (Native Forest, Exotic Tree Plantation-Forestry, Grassland, or Savanna), understory, tree and rock cover, distance to permanent water, and land-cover composition within 500-m and 5-km buffers (Supplementary Material Table S1). Land-cover classes and the buffer-composition metrics were derived from MapBiomas Uruguay (Collection 3; https://uruguay.mapbiomas.org). Camera models, sensor orientation, and the revision schedule are detailed in the Supplementary Material.

This study was observational: no animal was captured, handled, or manipulated at any stage, and camera-trap surveys of free-ranging wildlife require no mandatory wildlife-research permit in Uruguay. It followed an animal-use protocol approved by the national animal experimentation committee; fieldwork was conducted by personnel certified by that committee as animal experimenter and project coordinator. Owners or managers granted access to all sampled properties.

### Data preparation

Deer photographs were reviewed and tagged to species (*A. axis*, *S. gouazoubira*, or *O. bezoarticus*) in ExifPro (Kowalski 2013), and the associated date, time, and station metadata extracted with the camtrapR package (Niedballa et al. 2016) in R (R Core Team 2026). To avoid multiple counting of the same individual, photographs of a species taken within 1 h at a station were treated as a single independent detection unless different individuals could be distinguished in the images, which were recorded as separate detections; the 1-h threshold is widely used for mid- to large-sized mammals (Cravino and Brazeiro 2021; Cravino et al. 2023; Cravino and Brazeiro 2023). For activity analyses, each detection’s clock time was re-scaled to solar time (radians; sunrise ≈ π/2, sunset ≈ 3π/2) to compensate for seasonal variation in day length (Nouvellet et al. 2012; Frey et al. 2017; Vazquez et al. 2019), and, for nocturnal detections, lunar phase was obtained similarly (Kronfeld-Schor et al. 2013; Prugh and Golden 2014; Cravino and Brazeiro 2023); both were computed with the suncalc package (Thieurmel and Elmarhraoui 2019; Cravino and Brazeiro 2023).

### Single-species occupancy

We fitted Bayesian single-species occupancy models (Mackenzie and Royle 2005; Banks-Leite et al. 2014; Guillera-Arroita et al. 2014) with the PGOcc function of spOccupancy package (Doser et al. 2022), which uses Pólya–Gamma data augmentation (Polson et al. 2013). Each station’s detection history was a binary matrix over weekly occasions (with NA when the camera was inactive). Detection was modeled with four occasion-level covariates – days since the last maintenance, Julian day, within-occasion effort, and mean lunar phase – and the best detection submodel was selected by WAIC (the full four-covariate model was best supported for all three species and was retained; Supplementary Material Tables S4 and S5, Supplementary Material Figure S4). With detection so parameterized, we compared six occupancy submodels representing distinct ecological hypotheses – null, geographic (the invasion-axis gradient), local habitat, landscape composition at 500 m and at 5 km, and disturbance – by WAIC, and retained the best-supported model per species. We computed predicted occupancy (ψ) by habitat type (continuous covariates held at their means) and covariate-specific response curves. Covariate definitions (Supplementary Material Table S1), full model specifications, MCMC settings, and convergence diagnostics are given in the Supplementary Material. For *O. bezoarticus*, detected at only four of the seven stations within its endemic basaltic range and absent from the remaining 96 stations, we fitted an occupancy model restricted to that range. Detection was modeled with the same four occasion-level covariates used for the other species, but occupancy was kept as an intercept-only term given the small number of occupied sites, so that ψ is interpreted as a descriptive estimate of site use within the occupied range rather than as a habitat-association model.

### Two-species occupancy

To assess whether the two species of a dyad co-occur more or less often than expected under independence (MacKenzie et al. 2004), we used the conditional two-species co-occurrence model of Rota et al. (2016) (occuMulti, unmarked package; Fiske & Chandler (2011)), having found that multi-species community models (msPGOcc) failed to converge for our two-species datasets. This framework estimates a marginal occurrence term for each species and a second-order interaction term (f3): a negative f3 indicates co-occurrence below chance (spatial segregation), a positive f3 indicates positive association, and an interval spanning zero indicates independence. We focused on the *A. axis* / *S. gouazoubira* dyad (96 stations), comparing occupancy structures of increasing complexity by AIC while holding the interaction constant (f3 ∼ 1), and report the model with continuous habitat covariates as primary because the model with the categorical habitat factor suffered from separation (Supplementary Material Table S8); for the best model we computed each species’ conditional occupancy given the presence or absence of the other. For the *A. axis* / *O. bezoarticus* dyad, the co-occurrence subset comprised only the seven stations where *A. axis* was present, so its occupancy was effectively 1 and the interaction term was not estimable; spatial inference for that dyad therefore rests on the single-species models and the edaphogeographic separation between the species’ ranges. Finally, restricting the analysis to forest stations, we regressed understory cover on the relative abundance index (RAI; Sollmann et al. (2013)) of *A. axis* (linear and quadratic fits compared by AIC, with Spearman’s ρ as a distribution-free check) to test whether understory structure covaried with invader abundance; this analysis is correlational, and its interpretation is addressed in the Discussion. Pairwise correlations among site-level covariates are shown in Supplementary Material Figure S3.

### Diel activity and temporal overlap

We characterized the diel activity of each species and invasion level using kernel density estimation on solar-time distributions (function fitact, activity package; Rowcliffe (2019)) and classified each activity pattern following Azevedo et al. (2018) (classification thresholds in the Supplementary Material). Temporal overlap between species pairs in each context was quantified with the coefficient of overlap Δ (Ridout & Linkie (2009); overlap package), which ranges from 0 (no overlap) to 1 (identical activity); we applied the Δ1 estimator for samples of 20–75 detections and Δ4 for larger samples. Differences between circular distributions were tested with the Mardia–Watson–Wheeler test (circular package; Agostinelli & Lund (2022)), which returns a W statistic approximately distributed as χ². Both Δ and the Mardia–Watson–Wheeler test required at least 20 detections per group.

### Hierarchical temporal partitioning

To test whether the diel activity curve of one species shifts with the presence of the other, we applied the hierarchical modeling framework of Iannarilli et al. (2025). Each station × month constituted a sampling session, and the response was the binomial proportion of activity across bins of solar time, with, as the categorical predictor of interest, whether the other species was detected in that station-month (i.e., the contrast is between station-months with vs without a detection of the other species, not between invaded and uninvaded sites), and station as a random effect. For each focal species, we compared a trigonometric formulation and a hierarchical generalized additive model (HGAM, a cyclic spline over solar time; mgcv package; Wood (2017)) by AIC, and report the HGAM (Figure 6) because it consistently provided the better fit; the trigonometric–HGAM comparison is detailed in the Supplementary Material. To assess seasonal confounding, we refitted the HGAMs with season as an additional fixed effect (Supplementary Material Figure S2).

**Figure 5.**
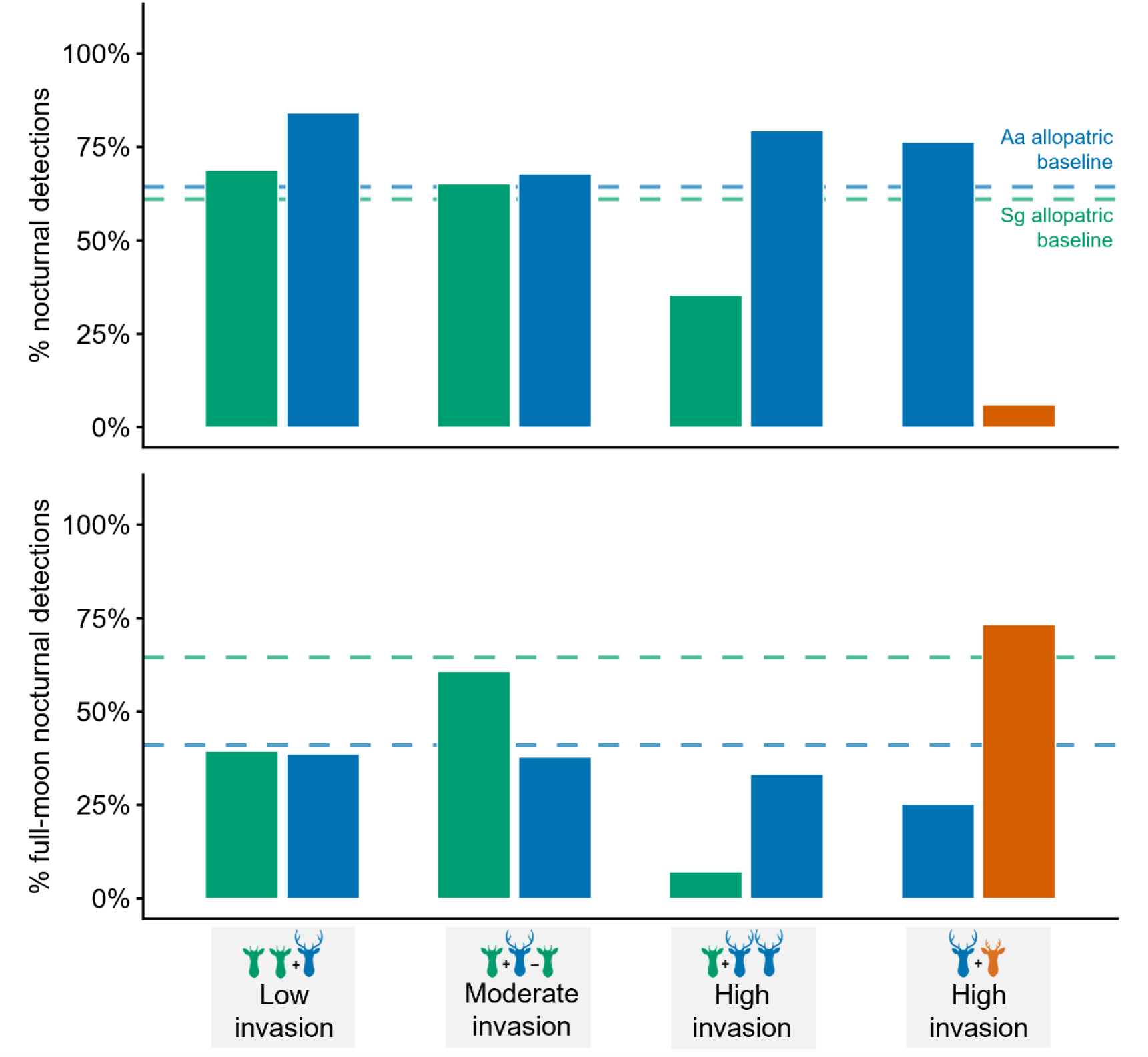
Nocturnal and lunar-phase activity of the three species across invasion levels. The figure summarises (a) the proportion of detections occurring at night and (b) the concentration of nocturnal detections within the lunar cycle (full-moon vs new-moon windows) for each species and context. Reference lines indicate the single-species allopatric baselines. Note the elevated full-moon activity of *S. gouazoubira* at moderate-invasion sites and no-invasion sites, and the suppression of moonlit activity of *S. gouazoubira* where *A. axis* numerically dominates (high-invasion sites). Species are identified by silhouette and color as in Figure 1. Alt text: Two stacked bar panels sharing a horizontal axis of four co-occurrence contexts. The upper panel gives the percentage of detections recorded at night, the lower the percentage of nocturnal detections falling around full moon. Bars are coloured by species and dashed horizontal lines mark each species’ baseline value where it occurs without the other. Both measures depart from those baselines as invasion advances, most markedly for *S. gouazoubira* at the highest invasion level.

**Figure 6.**
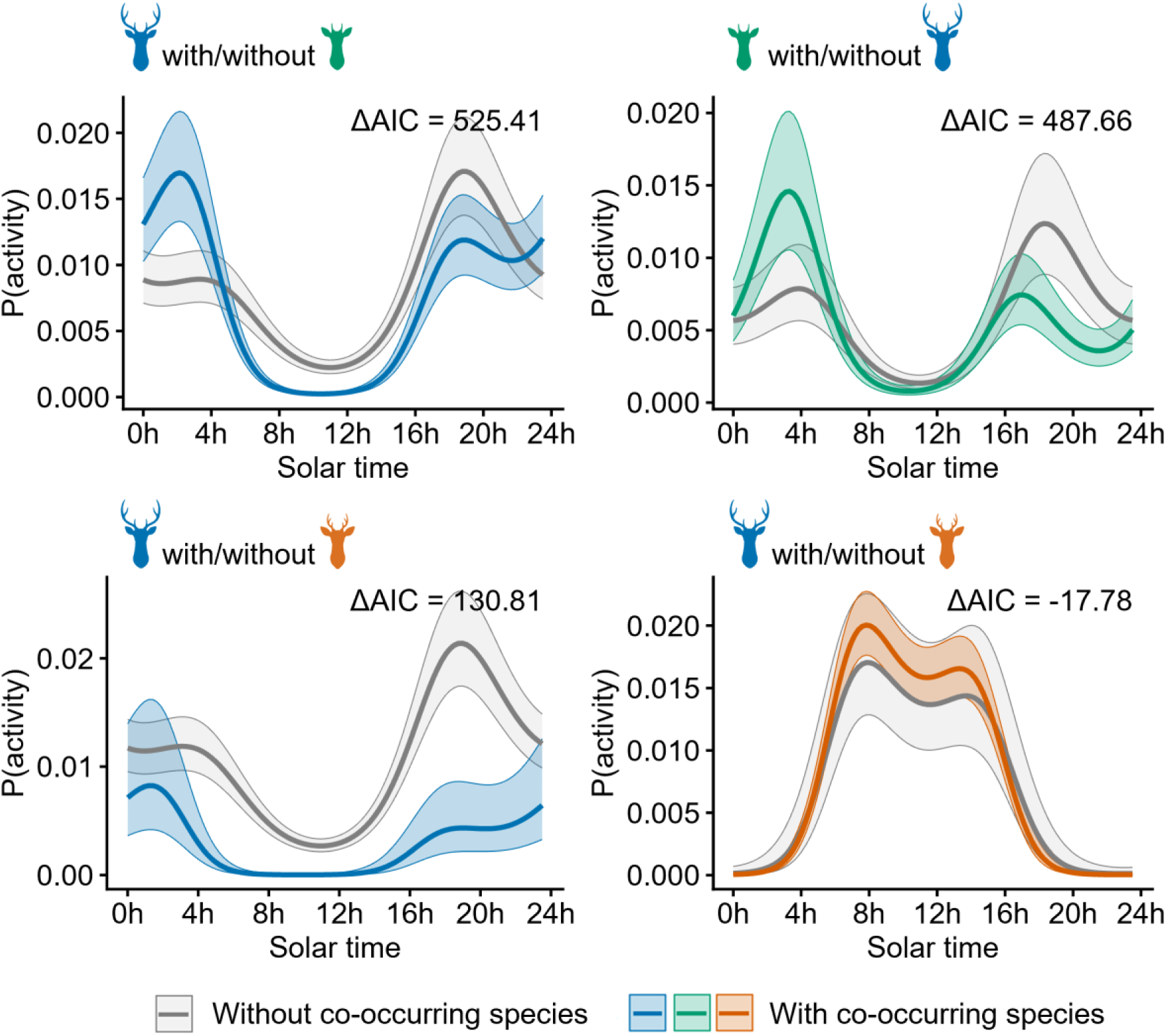
Hierarchical generalized additive models (HGAMs) of diel activity following Iannarilli et al. (2025), showing the model-predicted activity curve of each focal species with vs without the co-occurring species, for the four species-by-dyad combinations. Each panel shows the cyclic smooth of activity over solar time, fitted with and without the presence of the other species, with the ΔAIC favoring (or not) the inclusion of the co-occurring species. Three combinations show strong support for a diel shift in the presence of the other species (ΔAIC ≫ 0); the fourth – Ob with vs without Aa – shows a negative ΔAIC, indicating no support for a diel shift in Ob (serving as a comparative benchmark). Species are identified by silhouette and color as in Figure 1. Alt text: Four panels of modeled activity probability against solar time, one per focal species within its dyad. Each panel carries two curves with shaded 95% confidence ribbons, a grey curve for stations and months without the co-occurring species and a coloured curve for those with it, and the difference in AIC between the two models is printed in the panel. The curves separate clearly for *A. axis* and for *S. gouazoubira* in the invaded dyad and for *A. axis* alongside *O. bezoarticus*, but not for *O. bezoarticus* itself.

### Spatiotemporal avoidance and return-time dynamics

We tested for directional spatiotemporal avoidance between species with the permutation test of Niedballa et al. (2019) on inter-event intervals, complemented by linear mixed-effects models of log-transformed intervals (lme4 package; Bates et al. (2015)) with direction (A → B vs B → A) as the predictor of interest, station as a random intercept, and log sampling effort as a covariate. A direction coefficient departing from zero indicates that one species tends to follow the other more quickly, an asymmetry in how soon each species tends to follow the other at shared sites, which we interpret cautiously as directional spatiotemporal asymmetry. Increasingly complex models added invasion level, period of the day, camera-level overlap, and lunar phase, and were compared by AIC; negative-binomial GLMMs on transition counts (glmmTMB package, family nbinom2; Brooks et al. (2017)) and marginal-means contrasts (emmeans package; Lenth (2022)) served as complementary checks (Supplementary Material; Table S6). To exclude short-term avoidance of the human observer, we repeated the permutation tests after removing all detections within 7 days of a setup or revision event. For return-time dynamics within triples (A → B → A and B → A → B), we fitted analogous mixed models to log return time and compared species’ return rates using marginal means (Supplementary Material Table S7).

### Statistical Analyses

All analyses were conducted in R 4.6.0 (R Core Team 2026). Diel overlap and circular tests used the activity (Rowcliffe 2019), overlap (Ridout and Linkie 2009), and circular (Agostinelli and Lund 2022) packages; hierarchical activity models used mgcv (Wood 2017) and GLMMadaptive (Rizopoulos 2025); spatiotemporal-avoidance and return-time models used lme4 (Bates et al. 2015), glmmTMB (Brooks et al. 2017), and emmeans (Lenth 2022); and occupancy models used spOccupancy (Doser et al. 2022) and unmarked (Fiske and Chandler 2011). Model structures, formulas, and outputs are summarised per question in the analytical-framework table, with full specifications in the Supplementary Material. Species are abbreviated *A. axis* (Aa), *S. gouazoubira* (Sg), and *O. bezoarticus* (Ob) through the manuscript. We regarded an effect as statistically supported when its 95% confidence or credible interval excluded zero (α = 0.05); model selection used AIC (frequentist models) or WAIC (occupancy models), and Bayesian convergence was assessed with the Gelman–Rubin diagnostic (Rhat < 1.05; Gelman & Rubin (1992)). The R code and the analytical-framework table are provided in the Supplementary Material.

## Results

### Sampling effort

Over the eight-year monitoring period, the 103 camera-trap stations accumulated 154,590 camera–nights (Supplementary Material Table S3). *A. axis* was detected 25,623 independent times, *S. gouazoubira* 9,622 times, and *O. bezoarticus* 840 times. Relative abundance indices (RAI; detections per 100 camera–nights) for *A. axis* remained high throughout the period (ranging from 13.4 to 22.4 across years), while *S. gouazoubira* RAI peaked at 8.4 in 2021 and was lower, at 4.3–5.7 in 2023–2025. *O. bezoarticus* was detected with RAI values of 0.78–1.27, all from stations within the basaltic region. At the total-invasion sites, *S. gouazoubira* was not recorded across 80,239 camera-nights.

### Single-species occupancy and habitat selection

For all three species, the full detection model was best supported by WAIC (Aa, Sg, Ob all selected the four-covariate detection model; Supplementary Material Table S4). Detection probability increased with sampling effort within the occasion in all three species, and the effect of days since camera servicing differed in sign among them: positive in *O. bezoarticus* (β = +0.262) and, more weakly, in *A. axis* (β = +0.044), and negative in *S. gouazoubira* (β = -0.127, 95% CI [-0.176, -0.077]) – that is, detectability varied with the timing of human visits, in species-specific directions.

Habitat selection differed markedly between the two co-distributed species. For *A. axis*, the local-habitat model was best supported (WAIC = 18,395; ΔWAIC to next model = 36.2). *A. axis* was positively but imprecisely associated with savanna (β = +1.42, 95% CI [-0.71, 3.92]) and avoided both grassland (β = -3.91 [-5.61, -2.27]) and commercial forestry plantations (β = -1.82 [-3.32, -0.33]); it also avoided dense understories (β = - 1.03 [-1.79, -0.25]) and selected rocky substrates (β = +0.70 [0.08, 1.54]). For *S. gouazoubira*, the local-habitat model was likewise best supported (WAIC = 10,384; ΔWAIC = 18.5), but the species showed the opposite microhabitat profile: a strong positive association with understory cover (β = +1.93 [0.94, 3.06]) – the strongest single effect across all occupancy models – and with tree cover (β = +1.39 [0.49, 2.30]), with no selection for rocky substrate (β = -0.43, ns). Translating these coefficients into predicted occupancy by habitat type (continuous covariates held at their mean), *A. axis* reached its highest occupancy in savanna (ψ ≈ 0.95) and forest (ψ ≈ 0.83) and its lowest in grassland (ψ ≈ 0.09), whereas *S. gouazoubira* reached its highest occupancy in grassland and forestry (ψ ≈ 0.69 and 0.63) and forest (ψ ≈ 0.33), with low occupancy in savanna (ψ ≈ 0.14) (Figure 2; Supplementary Material Tables S12–S14; Supplementary Material Figure S5). Notably, both species occupied native forest with appreciable probability, indicating that forest functions as a shared habitat while the species diverge most strongly along the understory axis: *A. axis* avoids dense understories while *S. gouazoubira* requires it.

Within its endemic basaltic range, *O. bezoarticus* showed an appreciable but imprecisely estimated occupancy (ψ = 0.56, 95% CI 0.24–0.85; intercept-only model). We did not model its habitat associations: with detections at so few sites, a national-scale model would largely reproduce the species’ restricted geography through structural zeros rather than estimate habitat selection. The estimate therefore describes site use within the occupied range and indicates that *O. bezoarticus* is locally established there, consistent with a geographically restricted rather than a locally rare distribution.

### Understory structure and A. axis abundance

Because *S. gouazoubira* occupancy was strongly tied to understory cover, and because understory is the principal axis separating the two species, we examined whether understory structure itself covaried with *A. axis* abundance. Restricting the analysis to forest stations − thereby controlling for the large variation in understory among habitat types − understory cover declined steeply with *A. axis* relative abundance (linear regression β = -0.89 ± 0.17, t = -5.37, p < 0.001, R² = 0.39; Spearman’s ρ = -0.67, p < 0.001; a quadratic term did not improve the fit, indicating an approximately linear decline).

Predicted understory cover decreased from roughly 50 at forest sites without *A. axis* to roughly 13 at the highest observed *A. axis* relative abundance, a reduction of about 73% across the abundance gradient (Supplementary Material Figure S6). These measurements were taken at sites where *A. axis* was already established. We return in the Discussion to interpret this pattern and its limitations.

### Two-species occupancy and spatial association

The two-species co-occurrence model for the Aa/Sg dyad revealed strong spatial segregation (Figure 3). In the primary estimable model (continuous habitat covariates; the lower-AIC categorical-habitat model was non-identifiable through separation, Supplementary Material Table S8), the interaction term was negative and significant (f3 = -3.69, SE = 0.88, z = -4.19, p < 0.0001), and remained negative and significant in the simpler null-marginal model (f3 = -1.57, p < 0.001). Because this interaction is estimated after accounting for each species’ continuous habitat associations, the result indicates residual spatial segregation – negative co-occurrence beyond what the species’ divergent habitat associations alone would predict. We treat this as evidence of spatial segregation after accounting for the modeled covariates, rather than as direct evidence of competitive exclusion, since unmeasured geographic and habitat structure could also contribute; we return to the displacement interpretation in the synthesis below. Translating the interaction into conditional occupancy, the probability of *A. axis* occurrence fell from ψ ≈ 0.80 where *S. gouazoubira* was absent to ψ ≈ 0.21 where it was present (a 3.8-fold reduction), (Supplementary Material Table S15), while *S. gouazoubira* occurrence fell from ψ ≈ 0.67 to ψ ≈ 0.29 in the presence of *A. axis* (a 2.3-fold reduction).

For the Aa/Ob dyad, the co-occurrence subset comprised only stations where *A. axis* was present, so its occupancy was effectively 1 at all sites and the interaction term could not be meaningfully estimated (estimate unstable, SE > 100). Spatial inference for this dyad therefore rests on the single-species models – which place *O. bezoarticus* in open basaltic grasslands and *A. axis* across a broader range of habitats – and on the edaphogeographic separation between the species’ ranges, rather than on a direct estimate of spatial interaction.

### Diel activity and temporal overlap

Diel activity patterns differed strongly among species and across invasion levels. *A. axis* showed a predominantly cathemeral to mostly nocturnal pattern, depending on context (activity level ≈ 0.38–0.51 across contexts), with peaks around dusk and dawn. *S. gouazoubira* was also cathemeral, with a more uniform activity distribution and secondary crepuscular peaks (activity level ≈ 0.37–0.50). *O. bezoarticus*, in contrast, was predominantly diurnal (activity level 0.36), with detections concentrated between dawn and dusk and virtually no activity during the night. Temporal overlap (Δ) between *A. axis* and *S. gouazoubira* was very high when both species were measured in their respective single-species contexts (Δ = 0.955 when each species was measured where it occurred alone), but decreased markedly when both species co-occurred at the same sites. Overlap dropped to Δ = 0.451 at high-invasion sites (where Aa numerically dominated), to Δ = 0.679 at low-invasion sites, and remained high at Δ = 0.930 at moderate-invasion sites. In contrast, overlap between *A. axis* and *O. bezoarticus* was exceptionally low (Δ = 0.178), reflecting their fundamentally non-overlapping diel niches (Figure 4; Table 1).

**Table 1.**
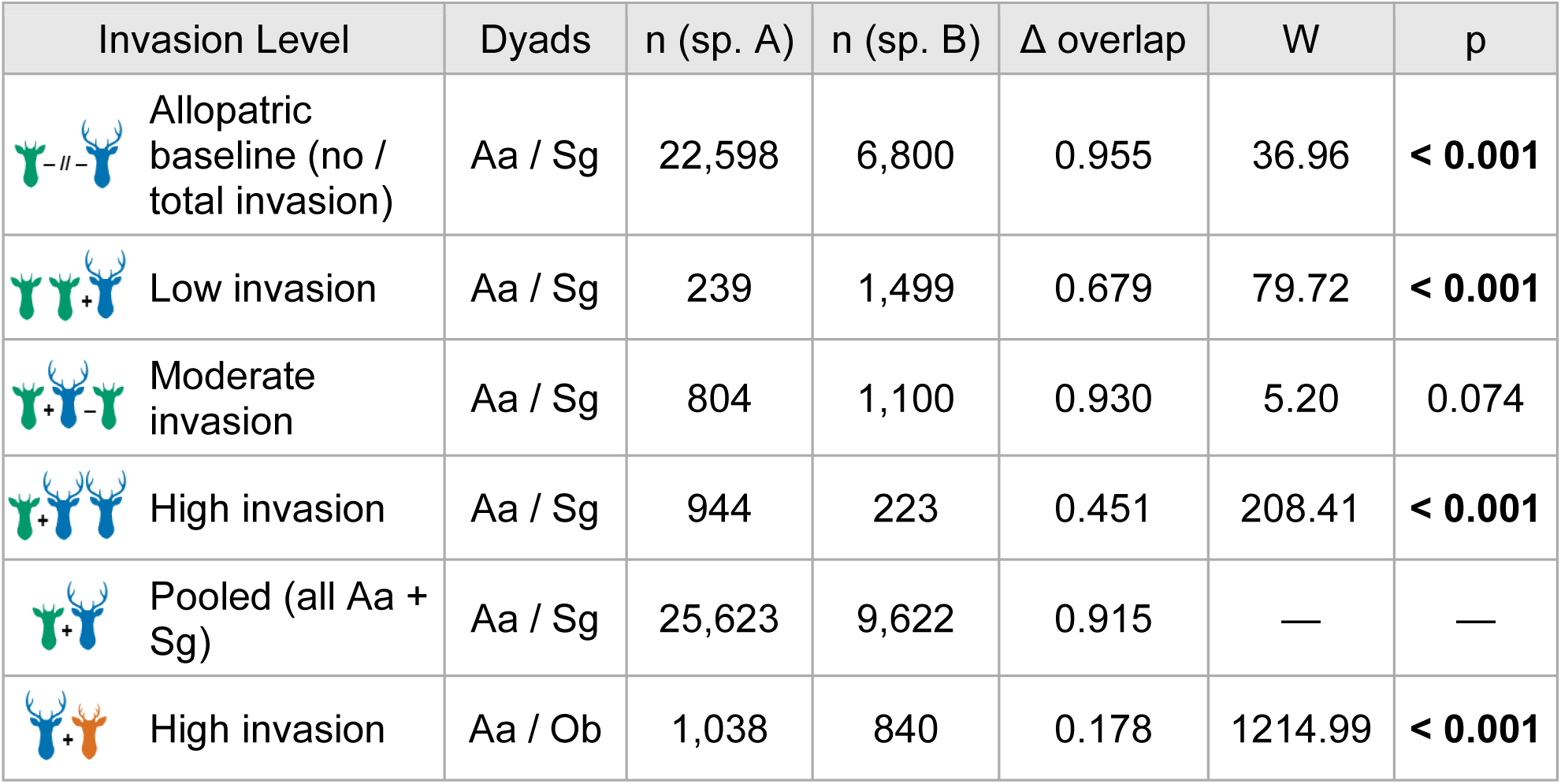
Diel activity overlaps and circular-distribution tests between species pairs across invasion levels. For each context and species pair the coefficient of overlap (Δ, from 0 = no overlap to 1 = identical activity) is reported with the number of detections contributing to each species’ activity curve (n), together with the Mardia–Watson–Wheeler test of the difference between the two circular (solar-time) distributions (W statistic, approximately χ²-distributed, with its p-value; larger W indicates more divergent diel distributions). The single-species row compares each species measured where it occurred alone; the pooled row uses all detections regardless of context (no circular test computed). The only non-significant W between *A. axis* and *S. gouazoubira* is at moderate-invasion sites, where the two species’ diel distributions converge.

| Invasion Level | Dyads | n (sp. A) | n (sp. B) | $\Delta$ overlap | W | p |
| --- | --- | --- | --- | --- | --- | --- |
| Allopatric baseline (no / total invasion) | Aa / Sg | 22,598 | 6,800 | 0.955 | 36.96 | **< 0.001** |
| Low invasion | Aa / Sg | 239 | 1,499 | 0.679 | 79.72 | **< 0.001** |
| Moderate invasion | Aa / Sg | 804 | 1,100 | 0.930 | 5.20 | 0.074 |
| High invasion | Aa / Sg | 944 | 223 | 0.451 | 208.41 | **< 0.001** |
| Pooled (all Aa + Sg) | Aa / Sg | 25,623 | 9,622 | 0.915 | — | — |
| High invasion | Aa / Ob | 1,038 | 840 | 0.178 | 1214.99 | **< 0.001** |

Mardia–Watson–Wheeler tests confirmed that the diel distributions of *A. axis* and *S. gouazoubira* differed significantly within most invasion levels (high-invasion sites: W = 208.4, p < 0.001; low-invasion sites: W = 79.7, p < 0.001; allopatric sites: W = 36.96, p < 0.001), with the notable exception of moderate-invasion sites, where the two species’ diel distributions did not differ statistically (W = 5.2, p = 0.074). This is the only context in which the two species showed no detectable divergence in diel activity – high overlap (Δ = 0.93) and statistically indistinguishable circular distributions – a pattern we interpret cautiously, as it may reflect recent contact at the front or lower statistical power there. The most extreme divergence was between *A. axis* and *O. bezoarticus* (W = 1215.0, p < 0.001), reflecting the diurnal–nocturnal segregation expected from their distinct ecologies.

### Lunar phase and nocturnality

Nocturnal activity varied substantially across species and contexts. *A. axis* was cathemeral to mostly nocturnal, depending on context, with 64–84% of detections occurring at night across contexts. *S. gouazoubira* showed generally lower levels of nocturnality (35–69%) but more variation, with nocturnality lowest at high-invasion sites (35%) and highest at moderate-invasion sites (65%). Most notably, *S. gouazoubira* detections in shared sites with *A. axis* tended to concentrate on the brighter portion of the lunar cycle. Among the lunar-active strata, *S. gouazoubira* had 64.5% of detections in the full-moon window at no-invasion sites (n = 2,874) and 60.8% at moderate-invasion sites (n = 497). *A. axis* at high-invasion sites showed a milder lunarphilic pattern (33.2%, n = 383), whereas *S. gouazoubira* at high-invasion sites – where *A. axis* dominated numerically – had only 7.1% in the full-moon window (n = 42). Given the small number of lunar-active detections at high-invasion sites (n = 42) relative to the other strata, we treat this apparent suppression of moonlit activity as suggestive rather than conclusive (Supplementary Material Table S9).

### Hierarchical temporal partitioning

The Iannarilli et al. (2025) hierarchical models revealed strong support for context-dependent diel shifts in three of the four species-by-dyad combinations and, critically, the absence of such shifts in the fourth. For the *Aa/Sg* dyad, both directions showed strong AIC support for including the co-occurring species in the model: *A. axis* with vs without *S. gouazoubira* yielded ΔAIC = 525.4 favoring the model with the other species, and *S. gouazoubira* with vs without *A. axis* yielded ΔAIC = 487.7 favoring the with-other model. For the *Aa/Ob* dyad, *A. axis* with vs without *O. bezoarticus* showed strong support for diel shift (ΔAIC = 130.8), but *O. bezoarticus* with vs without *A. axis* yielded ΔAIC = -17.8, meaning the null model was preferred by a substantial margin (Supplementary Material Table S11; Figure 6).

Three of the four panels show that diel activity does change in the presence of the other species – consistent with active context-dependent adjustment. But the fourth – *O. bezoarticus* with vs without *A. axis* – shows that the diel pattern of *O. bezoarticus* is essentially unchanged in station-months with vs without a detection of *A. axis*. The negative ΔAIC indicates no support for a detectable difference in the *O. bezoarticus* activity curve between station-months with and without an *A. axis* detection. We note, however, that the shared-site sample for *O. bezoarticus* is small, so this should be read as absence of a detectable response rather than as demonstrated invariance.

Taken together, the temporal and fine-scale spatiotemporal results place the two dyads at opposite ends of a partitioning gradient (Figure 9). The *A. axis* / *O. bezoarticus* dyad combines minimal diel overlap (Δ = 0.178) with pronounced directional avoidance, segregating along both the temporal and the fine-scale spatiotemporal axes. The *A. axis* / *S. gouazoubira* dyad, in contrast, overlaps strongly in activity time (Δ up to 0.955) and shows no directional avoidance in any context, indicating that these two species do not partition along either of these axes; their divergence emerging instead at the coarser scale of site occupancy and habitat described above.

### Spatiotemporal avoidance

Niedballa permutation tests on inter-event intervals revealed a striking contrast between the two dyads (Figure 7). For the *Aa/Sg* dyad, none of the three invasion levels showed statistically significant directional avoidance (high-invasion sites: p = 0.441; low-invasion sites: p = 0.667; moderate-invasion sites: p = 0.360). Linear mixed-effects models confirmed this, with the coefficient of direction (AB vs BA, log gap scale) close to zero and non-significant in all model variants (baseline: β = 0.085, 95% CI [-0.297, 0.467]; diel-corrected, delta-corrected, moon-controlled, and human-avoidance-controlled models all yielded |β| < 0.10 with CIs spanning zero; Supplementary Material Table S6). This pattern of statistical symmetry between AB and BA intervals indicates that *A. axis* and *S. gouazoubira* do not behaviorally avoid each other in time at the sites where they co-occur, despite the diel-overlap differences documented above.

**Figure 7.**
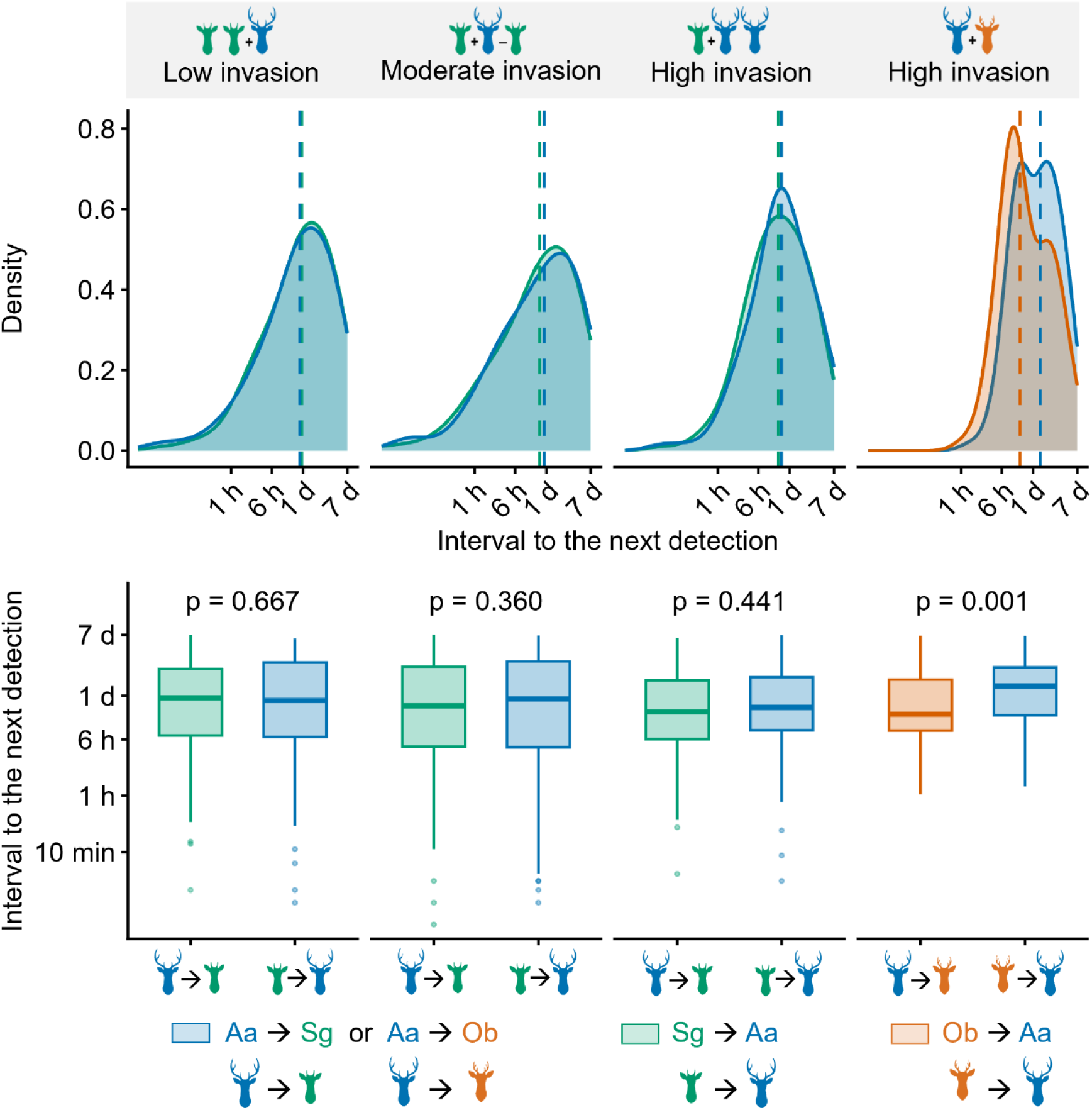
Fine-scale spatiotemporal avoidance between species pairs. Directional spatiotemporal avoidance following Niedballa et al. (2019): for each dyad, the distribution of time intervals between consecutive detections of different species in each direction (A→B vs B→A), with the corresponding permutation-test results; a symmetric distribution indicates no directional avoidance, whereas a shift between directions indicates that one species delays its use of a site after the other. The Aa/Sg dyad shows no directional avoidance in any context, whereas for the Aa/Ob dyad the interval from *A. axis* to *O. bezoarticus* is shorter than the reverse – that is, *O. bezoarticus* tends to follow *A. axis* sooner than *A. axis* follows *O. bezoarticus*, consistent with *A. axis* delaying its return to sites recently used by the native. Species are identified by silhouette and color as in Figure 1. Alt text: Two rows of four panels, one column per co-occurrence context. The upper row shows overlaid density curves of the interval to the next detection of the other species on a logarithmic time axis, with dashed vertical lines at the medians; the lower row shows the same intervals as box plots, one box per direction. Color marks the species arriving second, and the permutation p-value is printed above each pair of boxes. Only the *A. axis* and *O. bezoarticus* pair shows a marked difference between the two directions.

For the *Aa/Ob* dyad, in contrast, the permutation test showed strong directional avoidance (p < 0.001). The baseline mixed-effects model estimated a coefficient of direction (β_AB) = -0.444 (95% CI [-0.666, -0.223], p < 0.001), indicating that *O. bezoarticus* was typically detected soon after *A. axis*, whereas *A. axis* was markedly slower to reappear at sites recently used by *O. bezoarticus* (median Aa→Ob interval ≈ 800 min vs Ob→Aa ≈ 2,000 min). Read behaviorally, this longer delay before the invader returned to sites the native had just used – but not the reverse – is consistent with *A. axis*, rather than *O. bezoarticus*, adjusting its space use. Subsequent models that controlled for period of the day, lunar phase, and human-disturbance effects attenuated the estimate; its direction remained negative, but the confidence intervals included zero (controlled models: β_AB ≈ -0.28 to -0.29, Supplementary Material Table S6). The sensitivity analysis excluding detections within 7 days of camera maintenance confirmed the robustness of the avoidance pattern (sensitivity p-values: Aa/Sg contexts 0.400–0.609, Aa/Ob <0.001), demonstrating that the observed asymmetry is not an artifact of human disturbance.

### Return-time dynamics

Analysis of return times within triples (A → B → A and B → A → B sequences) provided additional information about post-encounter dynamics (Figure 8). For the *Aa/Sg* dyad, the baseline mixed-effects model on log return-time showed no significant difference between species (β = 0.22, 95% CI [-0.09, 0.53], p = 0.17), with median return times of 1,046– 1,322 minutes for *A. axis* and 813–1,336 minutes for *S. gouazoubira* across the three contexts (Supplementary Material Table S10). This symmetric pattern is consistent with the absence of directional avoidance in the pairs analysis.

**Figure 8.**
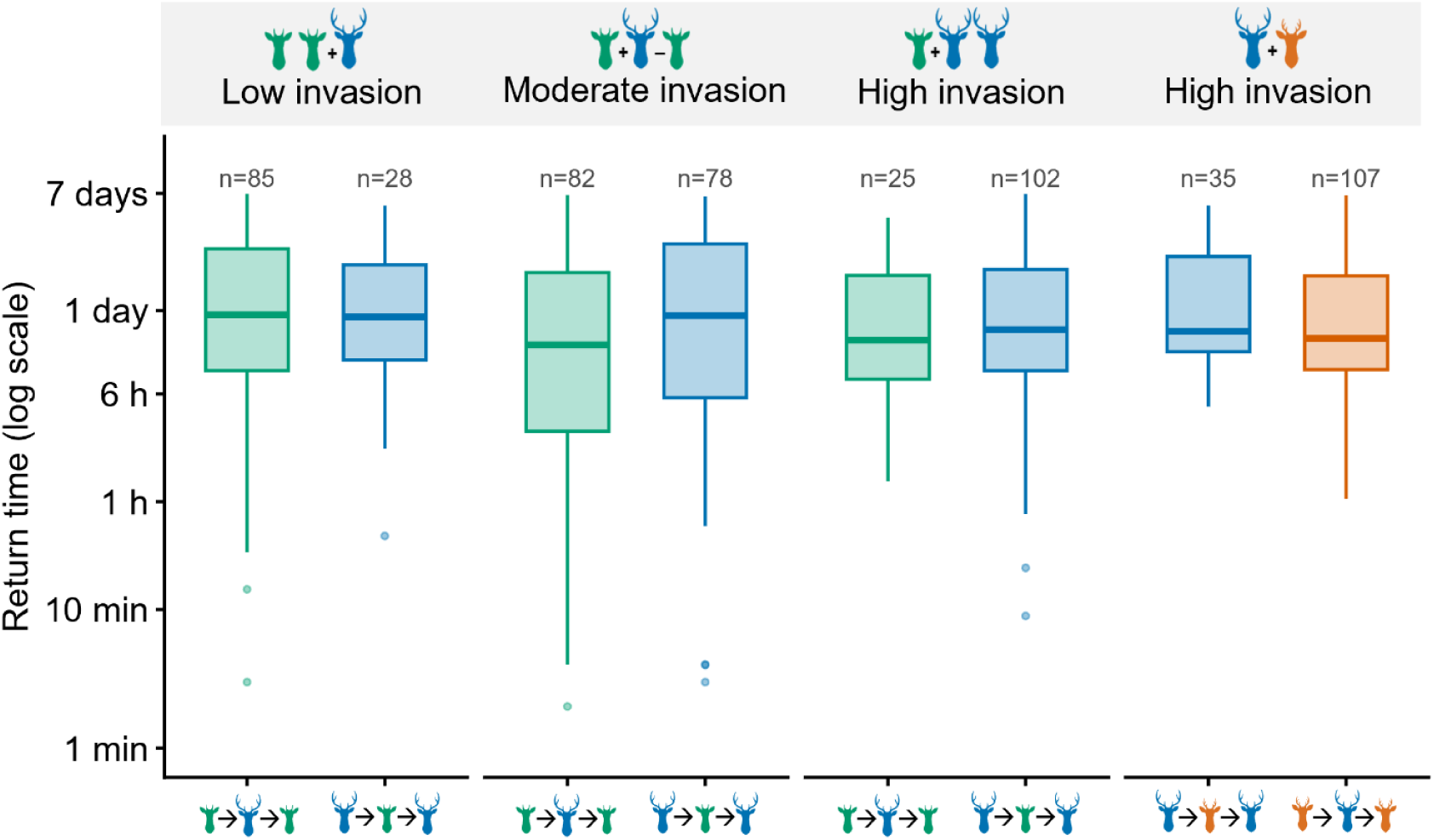
Return times within detection triples (A→B→A and B→A→B) across invasion levels, on a logarithmic scale; each box is the distribution of the time taken by the focal species to return to a site after the other intruded. Note that once an intrusion has occurred, return times are broadly similar between species. Species are identified by silhouette and color as in Figure 1. Alt text: Box plots of return time on a logarithmic axis, grouped into four panels by co-occurrence context, with one box per returning species and the number of sequences printed above it. Medians fall between roughly thirteen and twenty-two hours in every context, with no clear separation between the returning species.

**Figure 9.**
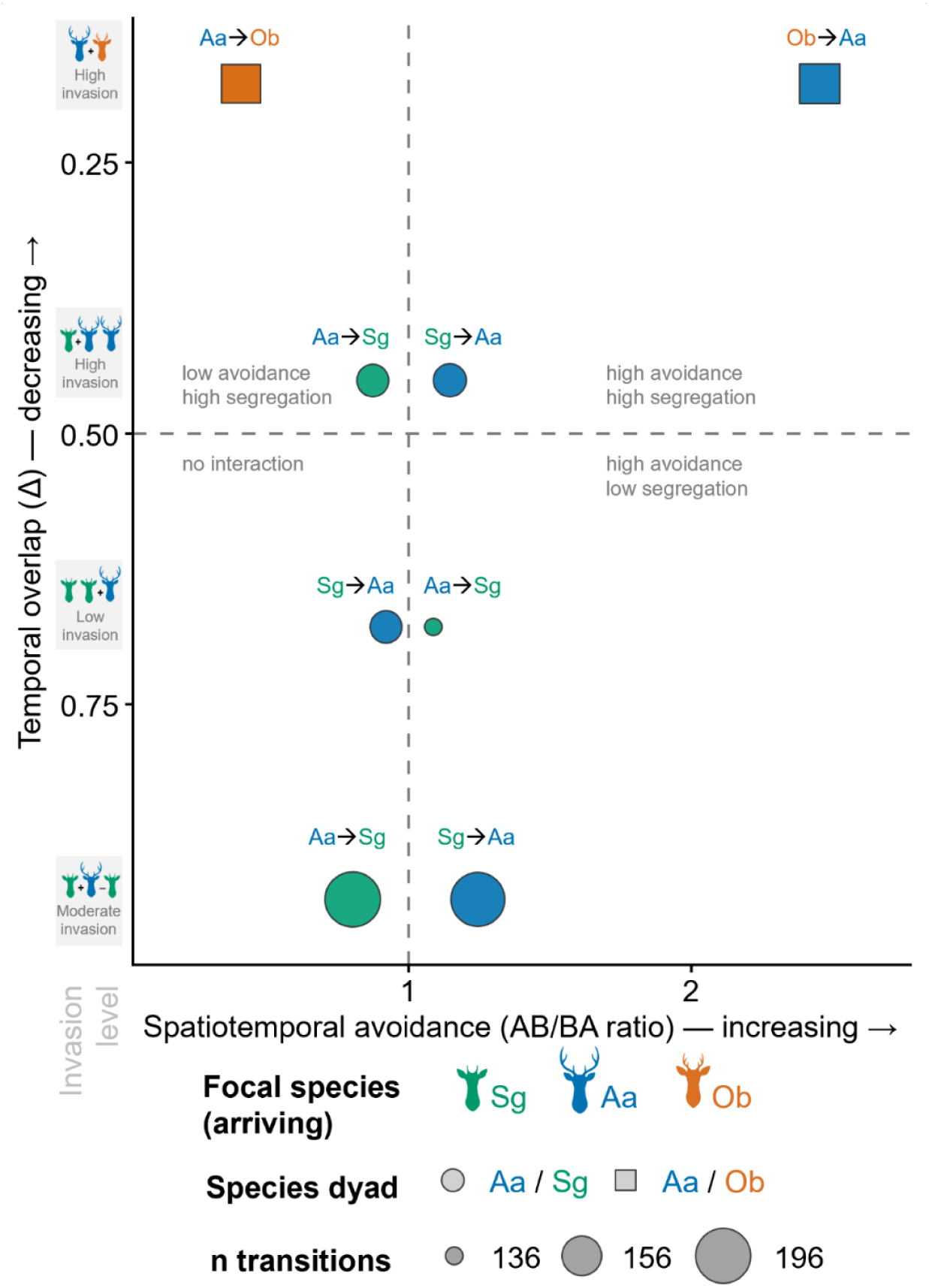
Synthesis of temporal and spatiotemporal niche partitioning across species pairs and invasion levels. Each point places a dyad × context combination according to its temporal overlap (Δ; vertical axis) and its degree of directional spatiotemporal avoidance (horizontal axis); symbol size is proportional to the number of directional transitions and color denotes the arriving species and symbol shape the species dyad. Combinations toward the lower left overlap strongly in activity time while showing little spatiotemporal avoidance, whereas those toward the upper right combine lower temporal overlap with stronger spatiotemporal avoidance. Species are identified by silhouette and color as in Figure 1. Alt text: Scatter plot with directional spatiotemporal avoidance on the horizontal axis and temporal overlap on the vertical axis, the vertical axis reversed so that temporal segregation increases upward. Dashed reference lines at an avoidance ratio of one and an overlap of 0.5 divide the plane into four labeled quadrants. Symbol shape marks the species dyad, fill color the arriving species and symbol size the number of transitions. The two *A. axis* and *O. bezoarticus* points sit at the top of the plot and far apart along the avoidance axis, whereas the *A. axis* and *S. gouazoubira* points cluster near the centre and the lower half.

For the *Aa/Ob* dyad, the baseline return-time model also showed no significant difference (β = 0.30, 95% CI [-0.10, 0.70], p = 0.13), with *A. axis* returning at a median of 1,016 min vs *O. bezoarticus* at 905 min. However, this similarity must be interpreted in light of the strong asymmetry in sample sizes: *A. axis* was the intervening (intruding) species in only 35 triples, whereas *O. bezoarticus* intervened between two *A. axis* detections in 107 triples – a disparity that largely reflects the species’ very different detection frequencies (a rare *O. bezoarticus* detection is far more often flanked by two *A. axis* detections than the reverse) rather than behaviour. The behavioral asymmetry between dyads, therefore, resides in the Niedballa pairs analysis and in the unequal number of detected sequences, not in the return-time durations themselves once an intrusion has already occurred.

### Change over time at the moderate-invasion level

Eight stations in Durazno department (moderate invasion site) operated continuously from March 2019 to December 2025 (19,880 camera-nights), allowing change to be assessed directly rather than inferred across sites. Over that period, *S. gouazoubira* declined from 16.3 to 0 (Figure 10) detections per 100 camera-nights, with the sharpest drop between 2020 and 2021 (13.7 to 4.4), and was not detected at all in 2025; *A. axis* increased from 3.7 to 5.2 over the same interval. Annual effort was near-constant (2,424– 2,920 camera-nights), and habitat covariates at these stations were unchanged (Supplementary material; Supplementary Material Table S16).

**Figure 10.**
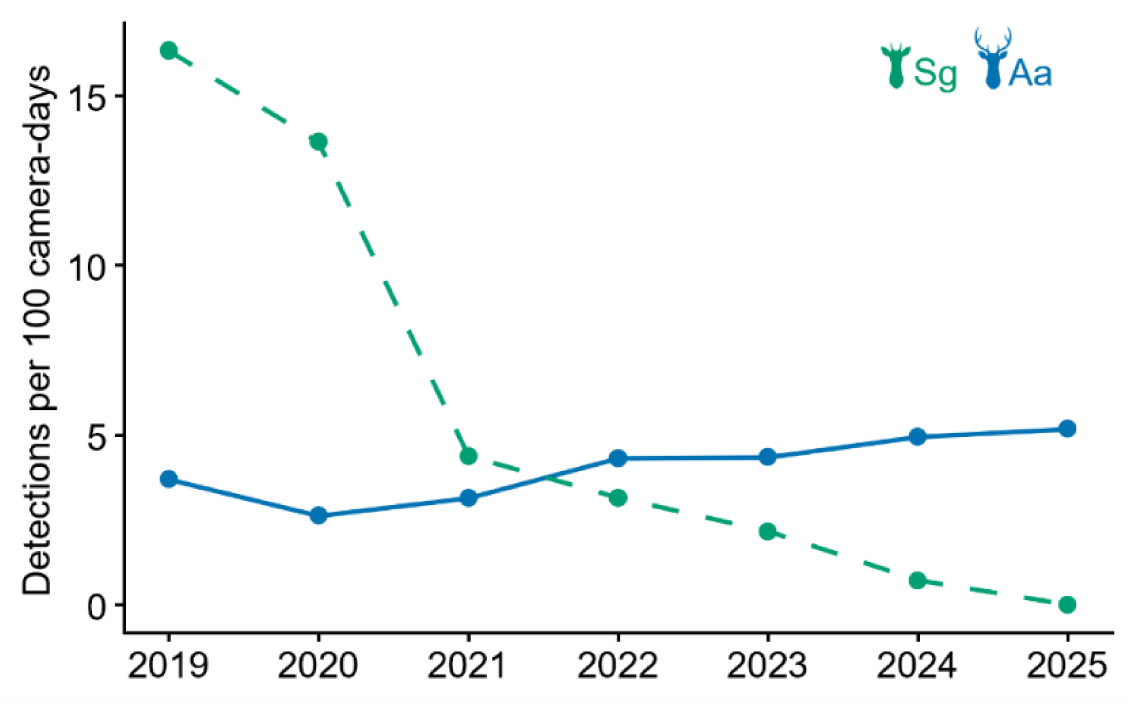
Seven-year trajectory of detection rates at the moderate-invasion site (eight stations in Durazno department monitored continuously, 2019–2025; 19,880 camera–nights). Points show annual detections per 100 camera–nights for *A. axis* (solid line) and *S. gouazoubira* (dashed line); effort is allocated by calendar day within each station’s deployment window (Supplementary Material Table S16). *S. gouazoubira* declined from 16.3 in 2019 to zero in 2025, with cameras operating throughout; *A. axis* rose from 3.7 to 5.2 over the same period. Species are identified by silhouette and color as in Figure 1. Alt text: Line chart of annual detection rates from 2019 to 2025 at eight camera stations. The line for *S. gouazoubira* falls steeply from about sixteen detections per 100 camera-nights to zero, while the line for *A. axis* rises gradually from under four to about five, the two crossing between 2021 and 2022.

## Discussion

Our eight-year camera-trap dataset shows that two native deer coexist with the same invader through fundamentally different mechanisms. At least for now, they do coexist, but this could change: the expansion of A. axis is ongoing and has not yet reached equilibrium in distribution and abundance relative to the availability of colonizable sites. *S. gouazoubira*, which shares much of its preferred habitat with *A. axis* (González and Martínez-Lanfranco 2010; Cravino and Brazeiro 2021; Cravino and Brazeiro 2023), shows a pattern of habitat-associated spatial segregation accompanied by context-dependent differences in diel activity: shifting diel overlap across invasion levels, strong spatial segregation at the site scale, and a tight dependence on the understory stratum that *A. axis* avoids and appears to degrade. *O. bezoarticus arerunguaensis*, by contrast, is segregated from *A. axis* along intrinsic axes – a predominantly diurnal activity period and an endemic distribution confined to the basaltic grasslands of north-central Uruguay (González et al. 2002; Cosse et al. 2009; González and Martínez-Lanfranco 2010; Cosse and González 2013; Gonzalez et al. 2023) – and we found no support for a detectable change in its behaviour in the presence of the invader. The contrast between the two dyads is the central result of this study: the same analytical framework that detects strong context-dependence in the Aa/Sg dyad reports its absence in the Aa/Ob dyad.

### Two coexistence mechanisms in a single landscape

Native species can persist alongside an invader either by adjusting their behavior and resource use, or because pre-existing niche differences limit overlap regardless of the invader (Schoener 1974; Kronfeld-Schor and Dayan 2003). These two routes are not mutually exclusive, but they may imply different futures: behavioral accommodation can, in principle, erode if the invader’s pressure intensifies, whereas segregation anchored in intrinsic niche differences is expected to be more stable (Chesson 2000). By studying two native cervids exposed to the same invader within the same national study system and analytical framework, we observed both routes simultaneously. The Aa/Ob dyad also serves as an internal control: when the hierarchical temporal models are applied to *O. bezoarticus,* they favor the null model of no diel shift, even though the same models detect large shifts for *A. axis* and *S. gouazoubira*. This guards the analysis against method-generated artifacts; the causal interpretation of the Aa/Sg dyad rests separately on the convergence of evidence set out below.

### The Aa/Sg dyad: habitat-mediated, context-dependent displacement

Four results point to the same interpretation for *S. gouazoubira.* First, although their preferred habitats do differ – *A. axis* reaches its highest predicted occupancy in savanna and avoids commercial plantations (β = -1.82 [-3.32, -0.33]), whereas *S. gouazoubira* occupies grassland and plantations – both species use native forest, with appreciable predicted occupancy there. Native forest is therefore the setting in which segregation can be assessed without differences in habitat preference alone accounting for it – consistent with the broad habitat use and ecological flexibility documented for the brown brocket across the Neotropics (Rivero et al. 2005; Rodrigues et al. 2014; Grotta-Neto et al. 2019; Grotta-Neto et al. 2024).

Second, within that shared habitat, the species diverge sharply along the understory axis: *S. gouazoubira* occupancy increases strongly with understory cover – the largest single effect in our occupancy analysis – whereas *A. axis* avoids dense understories and instead reaches its highest predicted occupancy in savanna, an open, parkland-like wooded savanna with a grassy ground layer and little woody understory. The relatively high predicted occupancy of *S. gouazoubira* in grassland should not be read as independence from woody cover: across the north-central and eastern regions of Uruguay, grasslands are typically interwoven with patches of hill forest and shrubby vegetation, so grassland stations frequently lie within reach of the wooded cover and understory on which the brocket depends. Consistent with this, the species’ strongest and most robust association was with the continuous understory and tree-cover gradients rather than with the categorical grassland class, and we accordingly base our habitat interpretation on the former. The same reasoning applies to plantations. Understory cover was recorded as absent at all 20 plantation stations, so brocket occupancy there (ψ ≈ 0.65) cannot rest on woody understory; what distinguishes the north-eastern stands where the species occurs from the western stands where it does not is the herbaceous ground layer (mean cover 36% against 6%) together with higher canopy cover (68% against 54%). Taken with the forest results, the brocket’s requirement is best described as cover at ground level, expressed as understory in native forest and as a herbaceous layer beneath plantation canopy. Neotropical cervids are comparatively tolerant of anthropogenic land cover in general (Costa et al. 2021); our results specify what that tolerance rests on here – not the origin of the canopy but the cover beneath it.

Third, the two-species model reveals strong spatial segregation that persists after controlling for these habitat associations: the interaction term is negative and significant, and the predicted occupancy of *S. gouazoubira* falls markedly from sites where *A. axis* is absent to sites where it is present. Crucially, this spatial segregation is not accompanied by fine-scale temporal avoidance: neither the permutation tests nor the return-time models detected directional avoidance in any context. Where the two species do co-occur, they do not stagger their visits in time. The displacement therefore operates at the level of site occupancy and habitat suitability, not through moment-to-moment temporal avoidance – a pattern paralleled in other cervid dyads that segregate spatially while overlapping strongly in time (cf. Chaudhary et al. 2020). In Argentine protected areas where it is established, *A. axis* occupies nearly all surveyed sites and overlaps in activity with most native mammals, although with displaced activity peaks (Shalom et al. 2025). Interference competition offers a plausible mechanism: axis deer are dominant over a native cervid at shared foraging sites, and the native’s selection for wooded habitat increased once axis deer were removed (Faas and Weckerly 2010), although that work was conducted at provisioned feeding sites.

Fourth, this sequence is not only spatial. At the moderate-invasion level, where the same eight stations operated continuously for seven years with near-constant effort, *S. gouazoubira* detection rates fell from 16.3 to 0 per 100 camera-nights while those of *A. axis* rose from 3.7 to 5.2, with local habitat unchanged. The cross-sectional gradient in invasion level therefore reflects a process that can be observed unfolding within a single locality, which is what licenses reading the levels as stages rather than as a static classification. Two qualifications apply. The decline was steepest between 2020 and 2021, a period coinciding with pandemic-related restrictions on movement and on hunting, which is the principal control on *A. axis* populations in Uruguay (Cravino et al. 2021); we cannot exclude a contribution from changing hunting pressure in those years, although *A. axis* detection rates at these stations did not exceed their 2019 level until 2022, and at the most heavily invaded sites they did not rise at all over the same interval. And because this is a single locality, the trajectory shows that displacement can occur, not that it will occur wherever the invader arrives.

Because the single-season occupancy models treat each site’s state as static over a multi-year sampling period, ψ is best interpreted as cumulative or long-term site use across the monitoring window rather than as a closed-population estimate of current occupancy at any single date. We therefore do not infer dynamic, moment-to-moment displacement from the occupancy estimates alone, but from their convergence with the activity and understory results.

The understory links these observations into a mechanism. *S. gouazoubira* is a browser that depends on dense understory for both forage and cover (Rivero et al. 2005; Andrade-Núñez and Mitchell Aide 2010; Black-Décima et al. 2010; Rodrigues et al. 2014; Nanni 2015; Rodrigues et al. 2017; Albanesi et al. 2019; Grotta-Neto et al. 2019; Weiler et al. 2020; Cravino and Brazeiro 2021; Cravino and Brazeiro 2023; Grotta-Neto et al. 2024; Martínez-Polanco 2026), and our analysis restricted to forest stations showed that understory cover declines steeply with *A. axis* abundance (a reduction of roughly 73% across the observed gradient). Together these results suggest that *A. axis* reduces the understory stratum on which *S. gouazoubira* depends, progressively rendering invaded sites less suitable for the native. This interpretation requires care: our understory measurements are cross-sectional and were taken where *A. axis* was already established, so we cannot fully separate active degradation by *A. axis* from a tendency of the invader to settle preferentially in forests that already had sparse understory. A partial exception is the moderate-invasion level, colonized by *A. axis* only recently: there, the understory has been exposed to the invader for far less time, so any pre-existing sparseness cannot be attributed to *A. axis*. This makes preferential settlement in already-degraded forest an unlikely sole explanation, consistent with degradation in progress, although we did not fit a front-restricted understory model and this space-for-time comparison rests on a single recently invaded context – we therefore regard it as suggestive rather than as direct evidence (Pickett 1989; Damgaard 2019). A dedicated longitudinal assessment of understory change at moderate-invasion sites is underway and will be reported elsewhere. Nonetheless, active habitat modification remains the most parsimonious hypothesis. Browsing and trampling by *A. axis*, and consequent reductions in understory vegetation and regeneration, have been documented in other parts of its introduced range (Nuñez et al. 2010; Mohanty et al. 2016; Cravino et al. 2021; Anujan et al. 2022), lending external plausibility to this mechanism.

The diel results add a temporal dimension. Activity overlap between the two species was very high when each was measured where it occurred alone (a comparison that necessarily contrasts sites as well as species), but lower where they co-occurred – most strikingly at high-invasion sites, where *A. axis* numerically dominates. The hierarchical models confirmed that each species’ diel curve shifts in the presence of the other. At the same time, moderate-invasion sites – where *S. gouazoubira* is declining relative to *A. axis* – were the only level at which the two species’ diel distributions were statistically indistinguishable, and overlap remained high. This is consistent with a front of recent contact where temporal partitioning has not yet developed, or where the remaining *S. gouazoubira* have not yet adjusted, in contrast to longer-established contexts where diel divergence is pronounced. An alternative reading follows from the local trajectory at this level: most *S. gouazoubira* detections there predate the species’ local collapse, so the pooled convergence may equally describe partitioning that ceased as the native faded rather than partitioning yet to develop. The two readings are not exclusive, and both mark this level as the stage at which the balance tipped. This behavioral reorganization is unlikely to reflect the invader alone: because *A. axis* is a game species in Uruguay – subject to regulated and often illegal hunting and highly attractive to poachers, in a country where poaching is a recognized threat to native fauna and enforcement is limited (Chouhy and Dabezies 2021; Dabezies et al. 2023; Dabezies et al. 2024) – its presence is associated with elevated hunter activity, poaching-related reports, and camera-trap theft at axis-occupied sites (Dabezies et al. 2023; Cravino 2024), so sites occupied by the invader may also expose the co-occurring native to greater human disturbance. A cover-dependent browser such as *S. gouazoubira* is expected to respond to this compound landscape of fear – the invader, hunting pressure, and reduced refuge – by contracting its spatial and temporal niche (Gaynor et al. 2019; Mendes et al. 2020; Gaynor et al. 2021; Hubbard 2021; Dammhahn et al. 2022; Palmer et al. 2022).

The lunar results point, more tentatively, in the same direction: *S. gouazoubira* concentrated its nocturnal activity around the full moon in most contexts, but this full-moon peak was markedly reduced at high-invasion sites. This is consistent with the abundant invader constraining the moonlit activity of the native, although the pattern rests on relatively few detections in that context and should be treated as suggestive (Kronfeld-Schor et al. 2013; Prugh and Golden 2014; Pratas-Santiago et al. 2017; Marques and Fabián 2018; Nekaris et al. 2021; Cravino and Brazeiro 2023). As with the diel shifts, these temporal patterns are best read as the finer-grained accompaniment to the dominant spatial and habitat signal, not as evidence of direct temporal avoidance, which our pairwise analyses did not support.

### The Aa/Ob dyad: intrinsic segregation with asymmetric behavioral avoidance

The picture for *O. bezoarticus* is qualitatively different. The two species’ diel niches barely overlap, and this separation is anchored in distinct circadian biology rather than in a response to one another: *O. bezoarticus* is predominantly diurnal while *A. axis* is cathemeral to mostly nocturnal, depending on context, a divergence consistent with the phylogenetic patterning of activity in Neotropical deer, in which Blastocerina tend to be diurnal and Odocoileina nocturnal (González and Martínez-Lanfranco 2010; Marques and Fabián 2018; Di Bitetti et al. 2020; Cravino and Brazeiro 2023; Grotta-Neto et al. 2024), and the hierarchical models show that the diel pattern of *O. bezoarticus* does not change in the presence of *A. axis*. The segregation also has a spatial and geological dimension:

*O. b. arerunguaensis* is endemic to the basaltic region of north-central Uruguay, where shallow basalt-derived soils support the extensive natural grasslands the subspecies requires (González et al. 2002; Cosse et al. 2009; González and Martínez-Lanfranco 2010; Gonzalez et al. 2023), whereas *A. axis* has expanded primarily through other soil and vegetation types (González and Martínez-Lanfranco 2010; Cravino et al. 2021; Cravino et al. 2023; Cravino and Brazeiro 2023). The two species are thus separated along an edaphogeographic axis that predates and is independent of their interaction. In this dyad, therefore, the dominant mechanism is intrinsic niche segregation – temporal, spatial and geological – rather than behavioral accommodation.

Against this backdrop, one behavioral signal did emerge, and it was asymmetric. The interval analysis pointed the same way: *A. axis* was slower to reappear at sites just used by *O. bezoarticus* than the reverse, and the hierarchical models showed the complementary pattern – *A. axis* shifted its diel activity in the presence of *O. bezoarticus*, whereas *O. bezoarticus* did not respond to *A. axis*. The sensitivity analysis excluding detections within seven days of camera maintenance confirmed that this signal is not an artefact of human disturbance. We interpret it cautiously, for two reasons. First, *A. axis* was far more abundant and was detected far more frequently than *O. bezoarticus*, so a shorter *A. axis* to *O. bezoarticus* interval may partly reflect the invader’s much higher baseline visitation rate rather than active tracking of the native; the permutation framework mitigates, but does not fully remove, this asymmetry in encounter rates (Dymit et al. 2025). Second, these analyses rest on the small number of sites shared within the subspecies’ restricted range, and once an intrusion had occurred, the two species’ return times did not differ – the asymmetry resides in the directional comparison and the unequal frequency of sequences, not in post-encounter behavior. With those caveats, the pattern remains noteworthy: to the extent that it reflects behavior, it is the invader that adjusts to the native and not the reverse – an unusual configuration relative to the cervid coexistence literature, where the native is typically the responding party (e.g. Garabedian et al. 2023). The two-species occupancy model was uninformative for this dyad because *A. axis* was present at all shared sites by construction, so spatial inference here rests on the single-species models and the edaphogeographic separation rather than on a direct co-occurrence estimate.

### Methodological considerations

Three methodological choices shaped our inferences and merit comment. First, for the temporal-partitioning analysis we relied on hierarchical generalized additive models rather than the trigonometric formulation. The two approaches agreed in direction in every case, but the HGAM consistently fit better (Supplementary Material Table S11) because it captures non-sinusoidal features of the activity curves – crepuscular peaks and asymmetric night-time activity – that a low-order trigonometric model smooths over. We therefore report the HGAM in the main text and retain the trigonometric results in the Supplementary Material for comparability with earlier studies. Second, for the two-species occupancy analysis we deliberately did not use the hierarchical community model (msPGOcc): with only two species, the community-level variance hyperparameters are poorly identified, and our attempts to fit that model produced unstable variance estimates. The conditional two-species formulation of Rota et al. (2016) sidesteps this problem by estimating the interaction term directly, and it converged cleanly. Within that framework, we report the model with continuous habitat covariates rather than the nominally lower-AIC model that included the categorical habitat factor, because the latter suffered from complete separation (occupancy probabilities driven to 0 or 1 in sparsely sampled habitat classes). Third, our analyses carry the limitations inherent to camera-trap observational data. Occupancy and co-occurrence models included site-level effects and habitat covariates, but cannot fully control for unmeasured within-site heterogeneity — for example, the structural differences between commercial plantations, which our single “Forestry” category collapses. This may be relevant because the north-eastern plantations, managed on longer rotations for solid wood, tend to develop more structural complexity and understory than the short-rotation pulpwood plantations of the west and center (DIEA-MGAP 2024). The understory–abundance analysis, as discussed above, is cross-sectional and cannot establish causality on its own, and the relative abundance index we used there is an imperfect proxy for true density that can be influenced by among-site detection differences (Sollmann et al. 2013; Palmer et al. 2018). Camera-trap theft and human interference were more frequent at axis-occupied sites; while consistent with elevated human activity there, this may also have reduced local sampling effort, an effect we accounted for by including effort as a covariate/offset. Detection itself carried this signal: detectability varied with the days since a camera was last visited, in directions that differed among species. Camera traps are far less invasive than capture, collaring or direct observation, but they are not invisible: the visits that keep them running are themselves detectable in the data, and our avoidance analyses were repeated excluding the week after each visit for precisely this reason. As camera traps spread far beyond academic use, into management, hunting and recreation, this deserves stating plainly: they should be deployed deliberately, and read with the awareness that the observer’s trace is part of what the camera records. These limitations temper specific interpretations but do not undermine the central comparative result, which rests on multiple convergent lines of evidence rather than on any single model.

More broadly, our results illustrate the value of exposing more than one native species to the same invader within a single system. Studied alone, either native would have supported a general claim that the other contradicts: that invasion drives behavioral displacement (from *S. gouazoubira*) or that it does not (from *O. bezoarticus*). Applying an identical analytical pipeline to both shows the contrast is ecological rather than methodological, and it offers a template for disentangling the heterogeneous, context-dependent effects of biological invasions on native communities (Ricciardi et al. 2013; Simberloff et al. 2013; Sapsford et al. 2020).

### Conclusions and conservation implications

Two native cervids exposed to the same invader in the same landscape did not respond in the same way. *S. gouazoubira* shares native forest with *A. axis* but is segregated from it in space and, within that shared habitat, along the understory stratum the brocket requires and the invader avoids; its diel activity shifts with the invader’s presence, and understory cover declines as *A. axis* becomes more abundant. *O. bezoarticus* is separated from *A. axis* by opposite diel niches and by the basaltic grasslands to which the subspecies is confined, and we found no support for a detectable behavioral response to the invader. Coexistence is achieved in both dyads, then, but through mechanisms that differ in kind: accommodation in one, pre-existing separation in the other. The contrast matters for conservation. In our analyses, the threatened pampas deer showed no detectable behavioral response to the invader, likely because its habitat and activity period differ strongly from those of *A. axis*, whereas for the locally threatened brown brocket we found no clear evidence of range or abundance reduction but did detect behavioral changes that could translate into population effects over the longer term. This is unlikely to reflect a general insensitivity of *O. bezoarticus* to introduced ungulates: in Argentina, pampas deer avoid paddocks at cattle stocking rates above 0.6 AU/ha and increase vigilance when feral pigs are nearby (Perez Carusi et al. 2017).

The two coexistence mechanisms imply different conservation trajectories. The coexistence of *S. gouazoubira* with *A. axis* appears conditional and potentially fragile: it depends on the availability of habitat with ground-level cover and limited invader pressure. Native forest is one such setting; commercial plantations are another, and our occupancy results place *S. gouazoubira* there at relatively high occupancy (ψ ≈ 0.65) in a habitat class the invader actively avoids. Brocket occupancy of plantations was concentrated in the north-eastern stands, which carry a developed herbaceous layer beneath the canopy, and these are also the stands from which *A. axis* is absent; because no station recorded both species in plantation, whether plantations can function as refugia under invasion pressure remains untested – a secondary conservation role for production forestry that warrants dedicated study (Simonetti et al. 2013; Rodrigues et al. 2014; Rodrigues et al. 2017; Iezzi et al. 2018; Iezzi et al. 2019; Ferreira et al. 2020; Iezzi et al. 2020). If, however, *A. axis* continues to spread and to reduce understory within these refugia, the spatial and habitat-based buffer that currently allows coexistence could erode. For *O. bezoarticus*, the intrinsic temporal and edaphogeographic segregation from *A. axis* suggests that direct competitive displacement by the invader is, for now, a lesser threat than for *S. gouazoubira*. The principal vulnerability of the endemic subspecies lies instead in its small and geographically restricted range: a population confined to the basaltic grasslands of north-central Uruguay is exposed to habitat conversion, fragmentation and stochastic risks regardless of the invader (González et al. 2002; Cosse et al. 2009; González and Martínez-Lanfranco 2010; Cosse and González 2013; Gonzalez et al. 2023). The asymmetric behavioral response we detected – *A. axis* adjusting to *O. bezoarticus* rather than the reverse – is, to our knowledge, an unusual configuration relative to the cervid coexistence literature, and, if it reflects behavior rather than differential abundance, it suggests that the consequences of *A. axis* presence for *O. bezoarticus* may be subtler than outright displacement. Monitoring the basaltic grassland populations as A. axis continues to expand will be essential to detect any change in this balance.

More generally, our results argue for treating behavior as a leading indicator. Distribution and abundance are lagging metrics: by the time a decline registers in them, the process is well advanced – at our moderate-invasion stations, diel convergence and a collapsing detection rate preceded complete non-detection in the final sampling year. Behavioral change is measurable years earlier, from the same camera-trap data already collected for monitoring, at no additional field cost. Where an invader is still spreading, as *A. axis* is, shifts in when and where natives are active may be the earliest warning conservation gets, and the cheapest to heed.

## Acknowledgments

We thank the many volunteers who accompanied us in the field, both in servicing camera-trap equipment and in reaching remote corners of the country; this study would not have been possible without their work.

We are grateful to the landowners and farm managers who granted access to their properties over the eight years of sampling and who supported the fieldwork as far as they were able: the Nogués, Acuña, Altuna, Signorelli and Arrarte families. We also thank PROBIDES, the Ministerio de Ambiente, Montes del Plata and LUMIN for granting access to the areas under their management and for logistical support during fieldwork.

The authors used a large language model (Claude, Anthropic) to assist with English-language editing of the manuscript and to review the analysis code. All data collection, statistical analyses, interpretations, and conclusions are the authors’ own.

## Data Availability

Camera-trap detection records, station-level covariates and sampling effort will be archived in a public repository on publication. Precise station coordinates are withheld because the study area includes the range of *O. bezoarticus arerunguaensis*, an endangered subspecies subject to illegal hunting. Station locations are reported at department level; exact coordinates are available from the corresponding author for research purposes.

