## Supplementary Material for "Living with the invader: two native deer, two outcomes along a gradient of axis deer invasion"

Supplementary Material for: Cravino Mol A, Mirazo S, Brazeiro A, Martínez-Lanfranco JA. Living with the invader: two native deer, two outcomes along a gradient of axis deer invasion. bioRxiv preprint. [Citation to be updated with journal, volume, pages and DOI upon publication.]

### ***Living with the invader: two native deer, two outcomes along a gradient of axis deer invasion***

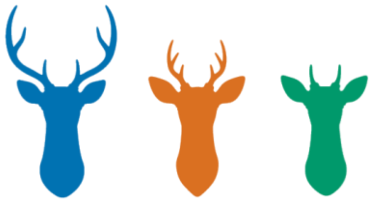

This document contains supplementary tables and figures supporting the main text. Contents: Table S1 (environmental covariates); Table S2 (camera-trap stations); Table S3 (relative abundance indices by year); Table S4 (detection model selection); Table S5 (detection coefficients); Table S6 (spatiotemporal-avoidance models); Table S7 (return-time models); Table S8 (two-species co-occurrence model selection and the discarded categorical model); Table S9 (nocturnal and lunar activity); Table S10 (return-time dynamics by dyad and invasion level); Table S11 (temporal-partitioning model selection); Tables S12–S14 (occupancy model selection, coefficients and predicted occupancy); Table S15 (two-species conditional occupancy); Table S16 (annual detections and effort at the moderate-invasion level, Durazno 2019–2025); Figure S1 (species distributions); Figure S2 (seasonal HGAMs); Figure S3 (covariate correlations); Figure S4 (detection probability); Figure S5 (occupancy responses); Figure S6 (understory vs *A. axis* abundance).

#### Analytical framework: questions, methods, data, and outputs

| Question | Analysis | Data used | Model / formula | R package | Main output |
|---|---|---|---|---|---|
| Q1. Diel activity & overlap by context | Diel solar overlap (Δ) + Mardia–Watson–Wheeler | All detections; solar-time radians | Kernel density (Dhat1/Dhat4); circular tests | activity, overlap, circular | Activity-level curves, Δ values, W statistics |
| Q2. Curve shifts with co-occurring species | Hierarchical GAM with cyclic cubic spline | Hourly counts × CT × month sessions | binomial(success, failure) ~ s(hour, bs='cc', by=oth_sp) + s(site, bs='re') | mgcv (bam), GLMMadaptive (mixed_model) | Marginal activity curves; ΔAIC vs the null model |
| Q3. Directional spatiotemporal avoidance | Niedballa permutation test + LMM | Pairs (A→B) and triples (A→B→A) with gap ≤ 7 days | Permutation; lmer(log_gap ~ direction × State + log_effort + (1\|CT)) | lme4, lmerTest, glmmTMB, emmeans, broom.mixed | Permutation p-values; direction coefficients; sensitivity analysis |
| Q4. Habitat selection (site & landscape) | Single-species occupancy (PGOcc) + WAIC | Detection history (J × K matrix, weekly occasions) | PGOcc(occ ~ habitat covs, det ~ days_CT_s + julian_day_s + effort_occ_s + moon_phase_s) | spOccupancy (PGOcc) | WAIC tables; best-model coefficients; predicted ψ |
| Q5. Joint spatial co-occurrence | Two-species co-occurrence model (Rota et al. 2016) | Two detection matrices (one per species) over weekly occasions | occuMulti(stateformulas = c(f1, f2, f3), detformulas = full p) | unmarked (occuMulti) | Interaction term f3 with 95% CI; conditional ψ given presence/absence of the other species |

*Note.* Q1–Q3 address the temporal and behavioral dimensions of coexistence; Q4–Q5 address the spatial dimension. All analyses were conducted in R version 4.6.0. Convergence and model-selection details for occupancy models are in Supplementary Material.

#### Supplementary Methods

This section provides the field protocol and full model specifications condensed from the main-text Methods; the analytical-framework table above summarises each analysis by question, and Table S1 defines the environmental covariates.

##### Field protocol

Cameras were Stealth Cam G42NG and GardePro NG units, attached to trees or wooden posts 30–50 cm above the ground with the sensor oriented horizontally toward an unobstructed view of the immediate surroundings. Whenever possible, cameras faced south and never east or west – and only northward when unavoidable – to prevent direct sunlight from reaching the sensor (the sun tracks across the northern sky in the Southern Hemisphere), thereby minimizing false triggers and overexposed images. Cameras were set to take three consecutive photographs per trigger, with sensor sensitivity set to high and no quiet period between triggers. No bait or scent attractants were used. Stations were revised at 3–10-month intervals (mean 175 days) to replace batteries and memory cards; revision intervals were progressively lengthened over the study as battery life and memory-card capacity under field conditions became better known, reducing the need for frequent visits.

##### Activity-pattern classification

Following Azevedo et al. (2018), activity patterns were classified from the proportion of records at night and twilight as: nocturnal (≥ 90% of records at night); mostly nocturnal (70–89% at night and twilight); crepuscular (≥ 80% of records around sunrise and sunset); cathemeral (30–69% at night and twilight); mostly diurnal (10–29% at night and twilight); and diurnal (active predominantly during daylight, ≥ 90% of records at daylight).

#### Hierarchical activity models

Following the framework of Iannarilli et al. (2025), two competing models were fitted per focal species. The trigonometric formulation used cosine and sine harmonics of 24-h and 12-h periods, each interacting with the presence/absence of the other species (oth_sp), fitted as a mixed-effects GLM (function mixed_model, GLMMadaptive; Rizopoulos 2025). The HGAM used a cyclic cubic spline for solar hour (k = 12) interacting with oth_sp, fitted with bam (mgcv; Wood 2017). Both used a binomial response with station as a random effect and were compared by AIC; the HGAM is reported in the main text. To assess seasonal confounding, the HGAMs were refitted with season (Spring, Summer, Autumn, Winter) as an additional fixed effect, deriving model-predicted diel curves for each season × oth_sp combination (Figure S2).

#### Spatiotemporal-avoidance and return-time models

Starting from the minimal model (direction + station random intercept + log effort), increasingly complex linear mixed-effects models added (i) invasion level, (ii) period of the day (is_night), (iii) camera-level temporal overlap (delta_CT), and (iv) lunar phase, and were compared by AIC (Table S6). As complementary checks we fitted negative-binomial GLMMs on directional transition counts (glmmTMB, family nbinom2; Brooks et al. 2017) and marginal-means contrasts (emmeans; Lenth 2022). For return-time dynamics within triples (A → B → A and B → A → B), we extracted the time between the intruder's detection and the focal species' return, fitted analogous linear mixed-effects models on the log return time, and compared species' return rates within each context by marginal means.

#### Occupancy model specifications

**Detection submodels.** For each species, six detection submodels were compared by WAIC: an intercept-only (null) model, each of the four covariates alone (days since maintenance, Julian day, within-occasion effort, mean lunar phase), and a full model with all four. The full model was best supported for all three species and retained (Table S4).

**Occupancy submodels.** The occupancy submodels were: null (intercept only); geographic, using the national west-to-east invasion-axis gradient within Uruguay as a broad descriptor of invasion geography rather than as evidence of a single continuous dispersal front; local habitat (habitat type, understory cover, tree cover, rock cover, distance to water); landscape composition at 500 m and, separately, at 5 km (proportion of forest, grassland, pasture, and wetland within the buffer); and disturbance (livestock pressure, human presence, exotic-species pressure, distance to water). All continuous covariates were standardised to mean 0 and SD 1 before fitting.

**Fitting.** Models were fitted with 10,000 posterior samples per chain across 3 chains, a 2,000-sample burn-in, and a thinning rate of 8 (final posterior = 3,000 samples). Convergence was assessed with the Gelman–Rubin diagnostic (Rhat), with all single-species models reaching Rhat < 1.05.

#### Two-species co-occurrence model

Multi-species community models (msPGOcc, spOccupancy) were initially fitted for consistency with the single-species framework but did not reach acceptable convergence for our two-species datasets, even after substantially increasing the number of posterior samples; we therefore adopted the conditional two-species model of Rota et al. (2016; occuMulti, unmarked). This model parameterises each joint occupancy state (neither species, only A, only B, or both) through natural parameters – marginal occurrence terms (f1, f2) and a second-order interaction term (f3). Model selection compared occupancy structures of increasing complexity by AIC (null marginals; continuous habitat covariates; disturbance covariates) while holding f3 ~ 1, with detection modeled by the full covariate set for each species. Because models including the categorical habitat factor (four levels) suffered from complete separation and non-identifiability (occupancy probabilities driven to 0 or 1 in sparsely sampled habitat classes), we report the model with continuous habitat covariates as the primary model; the model-selection table and the unstable categorical model are documented in Table S8.

#### Table S1. Environmental covariates

**Table S1.** Description of the local- and landscape-scale covariates recorded at each camera-trap station. Local covariates were assessed in the field within the camera's detection zone; landscape covariates were derived from land-cover layers within buffers of 500 m and 5 km around each station. All continuous covariates were standardized (mean 0, SD 1) before modelling.

| Covariate | Type | Description / units |
|---|---|---|
| Habitat | Categorical | Dominant habitat at the station: Forest, Forestry (commercial plantation), Grassland, or Savanna (wooded savanna). |
| Understory (Under) | Continuous | Field-estimated understory cover (%) within the detection zone. |
| Herbaceous cover (Herb) | Continuous | Field-estimated herbaceous ground cover (%) within the detection zone. |
| Tree cover (Tree) | Continuous | Field-estimated tree/canopy cover (%). |
| Rock cover (Rock) | Continuous | Field-estimated cover of rock/stony substrate (%). |
| Bare soil (BareSoil) | Continuous | Field-estimated cover of bare soil (%) within the detection zone. |
| Water distance (WaterDist) | Continuous | Distance to nearest permanent water (m). |
| Livestock | Continuous/index | Index of livestock presence at the station. |
| Human | Continuous/index | Index of human presence/activity at the station. |
| Dog | Continuous/index | Index of domestic dog presence. |
| EEI | Continuous/index | Index of exotic invasive species presence. |
| Forest / Grassland / Wetland / Pasture / Agriculture / Bare soil / Forestry / Eucalyptus / Pine / Other tree / Water (500 m, 5 km) | Continuous | Percent cover of each land-cover class within 500 m and 5 km buffers around the station. |
| Invasion axis | Continuous | Geographic gradient score summarising the broad west-to-east structure of *A. axis* invasion within Uruguay, used in the geographic occupancy model; this variable captures national invasion geography but is not interpreted as a single continuous dispersal front. |

#### Table S2. Camera-trap stations

**Table S2.** Camera-trap stations (n = 103) ordered by department and station code. For each station we report the department, dominant habitat, assigned invasion level, deployment period (setup to last retrieval) and cumulative sampling effort (camera-nights). Precise coordinates are not reported (see Data Availability in the main text). Invasion levels describe the local abundance of *A. axis* and follow the main text: no invasion (*A. axis* not detected); low, moderate and high invasion (*A. axis* present, in increasing relative abundance); and total invasion (*A. axis* detected and *S. gouazoubira* not detected). These levels are site-context classes within Uruguay that combine local invader abundance, invasion history and geography; they should not be read as independently replicated steps along a single national dispersal sequence. Stations in the basaltic range of *O. bezoarticus* fall within the high-invasion level and are identified as High invasion (Aa/Ob).

| Station | Department | Habitat | Context | Deployment | Camera-nights |
|---|---|---|---|---|---|
| ART1 | Artigas | Forest | Low invasion | Jan 2022–Jan 2023 | 363 |
| ART2 | Artigas | Forest | Low invasion | Jan 2022–Jan 2023 | 363 |
| ART3 | Artigas | Forest | Low invasion | Jan 2022–Jan 2023 | 363 |
| ART4 | Artigas | Forest | Low invasion | Jan 2022–Jan 2023 | 363 |
| ART5 | Artigas | Grassland | Low invasion | Jan 2022–Jan 2023 | 363 |
| ART6 | Artigas | Grassland | Low invasion | Jan 2022–Jan 2023 | 363 |
| CL1 | Cerro Largo | Forest | No invasion | Dec 2020–Jan 2023 | 737 |
| CL2 | Cerro Largo | Forest | No invasion | Dec 2020–Jan 2023 | 737 |
| CL3 | Cerro Largo | Forest | No invasion | Dec 2020–Jan 2023 | 737 |
| CL4 | Cerro Largo | Forest | No invasion | Dec 2020–Jan 2023 | 737 |
| CL5 | Cerro Largo | Forest | No invasion | Dec 2020–Jan 2023 | 737 |
| CL6 | Cerro Largo | Forest | No invasion | Dec 2020–Jan 2023 | 737 |
| CL7 | Cerro Largo | Forestry | No invasion | Dec 2020–Jan 2023 | 737 |
| CL8 | Cerro Largo | Forestry | No invasion | Dec 2020–Jan 2023 | 737 |
| COL1 | Colonia | Forest | Total invasion | Dec 2020–Jan 2026 | 1841 |
| COL2 | Colonia | Forest | Total invasion | Dec 2020–Jan 2026 | 1841 |
| COL3 | Colonia | Forest | Total invasion | Dec 2020–Jan 2026 | 1841 |
| COL4 | Colonia | Forest | Total invasion | Dec 2020–Jan 2026 | 1841 |
| COL5 | Colonia | Savanna | Total invasion | Dec 2020–Jan 2026 | 1841 |
| COL6 | Colonia | Savanna | Total invasion | Dec 2020–Jan 2026 | 1841 |
| COL7 | Colonia | Savanna | Total invasion | Dec 2020–Jan 2026 | 1841 |
| COL8 | Colonia | Savanna | Total invasion | Dec 2020–Jan 2026 | 1841 |
| DUR1 | Durazno | Forest | Moderate invasion | Mar 2019–Dec 2025 | 2485 |
| DUR2 | Durazno | Forest | Moderate invasion | Mar 2019–Dec 2025 | 2485 |

| Station | Department | Habitat | Context | Deployment | Camera-nights |
|---|---|---|---|---|---|
| DUR3 | Durazno | Forest | Moderate invasion | Mar 2019–Dec 2025 | 2485 |
| DUR4 | Durazno | Forest | Moderate invasion | Mar 2019–Dec 2025 | 2485 |
| DUR5 | Durazno | Forest | Moderate invasion | Mar 2019–Dec 2025 | 2485 |
| DUR6 | Durazno | Grassland | Moderate invasion | Mar 2019–Dec 2025 | 2485 |
| DUR7 | Durazno | Grassland | Moderate invasion | Mar 2019–Dec 2025 | 2485 |
| DUR8 | Durazno | Grassland | Moderate invasion | Mar 2019–Dec 2025 | 2485 |
| FL1 | Flores | Forest | Total invasion | Jan 2022–Dec 2024 | 1075 |
| FL10 | Flores | Forestry | Total invasion | Jan 2022–Dec 2024 | 1075 |
| FL11 | Flores | Grassland | Total invasion | Jan 2022–Dec 2024 | 1075 |
| FL12 | Flores | Grassland | Total invasion | Jan 2022–Dec 2024 | 1075 |
| FL2 | Flores | Forest | Total invasion | Jan 2022–Dec 2024 | 1075 |
| FL3 | Flores | Forest | Total invasion | Jan 2022–Dec 2024 | 1075 |
| FL4 | Flores | Savanna | Total invasion | Jan 2022–Dec 2024 | 1075 |
| FL5 | Flores | Savanna | Total invasion | Jan 2022–Dec 2024 | 1075 |
| FL6 | Flores | Savanna | Total invasion | Jan 2022–Dec 2024 | 1075 |
| FL7 | Flores | Forestry | Total invasion | Jan 2022–Dec 2024 | 1075 |
| FL8 | Flores | Forestry | Total invasion | Jan 2022–Dec 2024 | 1075 |
| FL9 | Flores | Forestry | Total invasion | Jan 2022–Dec 2024 | 1075 |
| PAY1 | Paysandu | Forest | Total invasion | Jan 2018–Jan 2026 | 2919 |
| PAY10 | Paysandu | Forestry | Total invasion | Jan 2018–Jan 2026 | 2919 |
| PAY11 | Paysandu | Grassland | Total invasion | Jan 2018–Jan 2026 | 2919 |
| PAY12 | Paysandu | Grassland | Total invasion | Jan 2018–Jan 2026 | 2919 |
| PAY13 | Paysandu | Grassland | Total invasion | Jan 2018–Jan 2026 | 2919 |
| PAY2 | Paysandu | Forest | Total invasion | Jan 2018–Jan 2026 | 2919 |
| PAY3 | Paysandu | Forest | Total invasion | Jan 2018–Jan 2026 | 2919 |
| PAY4 | Paysandu | Savanna | Total invasion | Jan 2018–Jan 2026 | 2919 |
| PAY5 | Paysandu | Savanna | Total invasion | Jan 2018–Jan 2026 | 2919 |
| PAY6 | Paysandu | Savanna | Total invasion | Jan 2018–Jan 2026 | 2919 |
| PAY7 | Paysandu | Forestry | Total invasion | Jan 2018–Jan 2026 | 2919 |
| PAY8 | Paysandu | Forestry | Total invasion | Jan 2018–Jan 2026 | 2919 |
| PAY9 | Paysandu | Forestry | Total invasion | Jan 2018–Jan 2026 | 2919 |
| RN1 | Rio Negro | Forest | Total invasion | Dec 2020–Dec 2025 | 1833 |
| RN2 | Rio Negro | Forest | Total invasion | Dec 2020–Dec 2025 | 1833 |
| RN3 | Rio Negro | Forest | Total invasion | Dec 2020–Dec 2025 | 1833 |

| Station | Department | Habitat | Context | Deployment | Camera-nights |
|---|---|---|---|---|---|
| RN4 | Rio Negro | Savanna | Total invasion | Dec 2020–Dec 2025 | 1833 |
| RN5 | Rio Negro | Savanna | Total invasion | Dec 2020–Dec 2025 | 1833 |
| RN6 | Rio Negro | Savanna | Total invasion | Dec 2020–Dec 2025 | 1833 |
| RN7 | Rio Negro | Forestry | Total invasion | Dec 2020–Dec 2025 | 1833 |
| RN8 | Rio Negro | Forestry | Total invasion | Dec 2020–Dec 2025 | 1833 |
| ROC1 | Rocha | Grassland | Low invasion | Feb 2020–Mar 2023 | 1126 |
| ROC10 | Rocha | Forest | High invasion | Feb 2020–Mar 2023 | 1126 |
| ROC11 | Rocha | Forest | High invasion | Feb 2020–Mar 2023 | 1126 |
| ROC12 | Rocha | Forest | High invasion | Feb 2020–Mar 2023 | 1126 |
| ROC13 | Rocha | Grassland | High invasion | Feb 2020–Mar 2023 | 1126 |
| ROC14 | Rocha | Grassland | High invasion | Feb 2020–Mar 2023 | 1126 |
| ROC15 | Rocha | Grassland | High invasion | Feb 2020–Mar 2023 | 1126 |
| ROC2 | Rocha | Grassland | Low invasion | Feb 2020–Mar 2023 | 1126 |
| ROC3 | Rocha | Grassland | Low invasion | Feb 2020–Mar 2023 | 1126 |
| ROC4 | Rocha | Forest | Low invasion | Feb 2020–Mar 2023 | 1126 |
| ROC5 | Rocha | Forest | Low invasion | Feb 2020–Mar 2023 | 1126 |
| ROC6 | Rocha | Forest | Low invasion | Feb 2020–Mar 2023 | 1126 |
| ROC7 | Rocha | Forest | Low invasion | Feb 2020–Mar 2023 | 1126 |
| ROC8 | Rocha | Forest | High invasion | Feb 2020–Mar 2023 | 1126 |
| ROC9 | Rocha | Forest | High invasion | Feb 2020–Mar 2023 | 1126 |
| SAL1 | Salto | Forest | High invasion (Aa/Ob) | Jan 2022–Jan 2025 | 1091 |
| SAL2 | Salto | Forest | High invasion (Aa/Ob) | Jan 2022–Jan 2025 | 1091 |
| SAL3 | Salto | Forest | High invasion (Aa/Ob) | Jan 2022–Jan 2025 | 1091 |
| SAL4 | Salto | Grassland | High invasion (Aa/Ob) | Jan 2022–Jan 2025 | 1091 |
| SAL5 | Salto | Grassland | High invasion (Aa/Ob) | Jan 2022–Jan 2025 | 1091 |
| SAL6 | Salto | Grassland | High invasion (Aa/Ob) | Jan 2022–Jan 2025 | 1091 |
| SAL7 | Salto | Grassland | High invasion (Aa/Ob) | Jan 2022–Jan 2025 | 1091 |
| TAC1 | Tacuarembo | Forest | No invasion | Dec 2021–Jan 2026 | 1477 |
| TAC10 | Tacuarembo | Forestry | No invasion | Dec 2021–Jan 2026 | 1477 |
| TAC11 | Tacuarembo | Grassland | No invasion | Dec 2021–Jan 2026 | 1477 |
| TAC12 | Tacuarembo | Grassland | No invasion | Dec 2021–Jan 2026 | 1477 |
| TAC2 | Tacuarembo | Forest | No invasion | Dec 2021–Jan 2026 | 1477 |
| TAC3 | Tacuarembo | Forest | No invasion | Dec 2021–Jan 2026 | 1477 |
| TAC4 | Tacuarembo | Forest | No invasion | Dec 2021–Jan 2026 | 1477 |

| Station | Department | Habitat | Context | Deployment | Camera-nights |
|---|---|---|---|---|---|
| TAC5 | Tacuarembo | Forest | No invasion | Dec 2021–Jan 2026 | 1477 |
| TAC6 | Tacuarembo | Forestry | No invasion | Dec 2021–Jan 2026 | 1477 |
| TAC7 | Tacuarembo | Forestry | No invasion | Dec 2021–Jan 2026 | 1477 |
| TAC8 | Tacuarembo | Forestry | No invasion | Dec 2021–Jan 2026 | 1477 |
| TAC9 | Tacuarembo | Forestry | No invasion | Dec 2021–Jan 2026 | 1477 |
| TT1 | Treinta y Tres | Forest | No invasion | Feb 2021–Dec 2022 | 691 |
| TT2 | Treinta y Tres | Forest | No invasion | Feb 2021–Dec 2022 | 691 |
| TT3 | Treinta y Tres | Forest | No invasion | Feb 2021–Dec 2022 | 691 |
| TT4 | Treinta y Tres | Forestry | No invasion | Feb 2021–Dec 2022 | 691 |
| TT5 | Treinta y Tres | Forestry | No invasion | Feb 2021–Dec 2022 | 691 |
| TT6 | Treinta y Tres | Forestry | No invasion | Feb 2021–Dec 2022 | 691 |

*Note.* Total cumulative sampling effort across all stations = 154,590 camera-nights. Deployment periods span January 2018 to January 2026; individual stations were active for between ~1 and ~8 years (363–2,919 days) depending on access, theft and equipment availability.

#### Table S3. Relative abundance indices by year

**Table S3.** Relative abundance index (RAI; detections per 100 camera-nights) for each species by year, with the number of detections and the sampling effort (camera-nights). RAI provides a coarse index of relative abundance and is not a density estimate. *O. bezoarticus* was detected only in 2022–2024 and only at stations within its basaltic-grassland range. The 2025 effort includes the final camera-retrieval period through January 2026, during which no focal-species detections were recorded; per-year sampling effort therefore sums to the study total of 154,590 camera-nights.

| Species | Year | Detections | Effort (camera-nights) | RAI |
|---|---|---|---|---|
| *A. axis* | 2018 | 775 | 4,628 | 16.75 |
| *A. axis* | 2019 | 1,606 | 7,177 | 22.38 |
| *A. axis* | 2020 | 1,927 | 12,497 | 15.42 |
| *A. axis* | 2021 | 3,474 | 23,994 | 14.48 |
| *A. axis* | 2022 | 5,024 | 37,509 | 13.39 |
| *A. axis* | 2023 | 5,291 | 26,192 | 20.20 |
| *A. axis* | 2024 | 5,056 | 24,648 | 20.51 |
| *A. axis* | 2025 | 2,470 | 17,945 | 13.76 |
| *S. gouazoubira* | 2019 | 397 | 7,177 | 5.53 |
| *S. gouazoubira* | 2020 | 936 | 12,497 | 7.49 |
| *S. gouazoubira* | 2021 | 2,024 | 23,994 | 8.44 |
| *S. gouazoubira* | 2022 | 2,928 | 37,509 | 7.81 |
| *S. gouazoubira* | 2023 | 1,138 | 26,192 | 4.34 |
| *S. gouazoubira* | 2024 | 1,184 | 24,648 | 4.80 |
| *S. gouazoubira* | 2025 | 1,015 | 17,945 | 5.66 |
| *O. bezoarticus* | 2022 | 294 | 37,509 | 0.78 |
| *O. bezoarticus* | 2023 | 333 | 26,192 | 1.27 |
| *O. bezoarticus* | 2024 | 213 | 24,648 | 0.86 |

#### Table S4. Detection model selection

**Table S4.** Model selection (WAIC) for the detection submodels of each species, fitted before the occupancy structure. Candidate models: null (intercept only), single-covariate models (days since maintenance, Julian day, within-occasion effort, lunar phase), and the full model with all four covariates. For all three species the full detection model was best supported and was retained for the occupancy analyses. ΔWAIC is the difference from the best model.

| Species | Detection model | WAIC | ΔWAIC |
|---|---|---|---|
| *A. axis* | full (4 covariates) | 18,433.0 | 0.0 |
| *A. axis* | effort only | 18,436.9 | 4.0 |
| *A. axis* | days since maint. only | 18,550.9 | 117.9 |
| *A. axis* | Julian day only | 18,551.3 | 118.3 |
| *A. axis* | null | 18,552.5 | 119.6 |
| *A. axis* | lunar phase only | 18,553.6 | 120.7 |
| *S. gouazoubira* | full (4 covariates) | 10,435.9 | 0.0 |
| *S. gouazoubira* | effort only | 10,464.6 | 28.7 |
| *S. gouazoubira* | days since maint. only | 10,510.5 | 74.6 |
| *S. gouazoubira* | Julian day only | 10,526.5 | 90.6 |
| *S. gouazoubira* | null | 10,535.7 | 99.8 |
| *S. gouazoubira* | lunar phase only | 10,535.8 | 99.9 |
| *O. bezoarticus* | full (4 covariates) | 860.2 | 0.0 |
| *O. bezoarticus* | effort only | 865.4 | 5.2 |
| *O. bezoarticus* | days since maint. only | 868.5 | 8.3 |
| *O. bezoarticus* | null | 869.8 | 9.6 |
| *O. bezoarticus* | lunar phase only | 871.1 | 10.9 |
| *O. bezoarticus* | Julian day only | 871.7 | 11.5 |

#### Table S5. Detection coefficients

**Table S5.** Detection coefficients (posterior median and 95% credible interval, logit scale) from the full detection model for each species. Coefficients whose interval excludes zero are in bold. Note that the effect of days since maintenance differs in sign among species: positive in *O. bezoarticus* and, more weakly, in *A. axis*, and negative in *S. gouazoubira*; and the consistently positive effect of within-occasion effort in all species.

| Species | Covariate | Estimate | 95% CI |
|---|---|---|---|
| *A. axis* | Intercept | 0.216 | **[0.168, 0.255]** |
| *A. axis* | days since maint. | 0.044 | **[0.011, 0.078]** |
| *A. axis* | Julian day | 0.026 | [-0.007, 0.061] |
| *A. axis* | effort | 0.635 | **[0.377, 1.091]** |
| *A. axis* | lunar phase | -0.014 | [-0.049, 0.020] |
| *S. gouazoubira* | Intercept | -0.215 | **[-0.265, -0.165]** |
| *S. gouazoubira* | days since maint. | -0.127 | **[-0.176, -0.077]** |
| *S. gouazoubira* | Julian day | 0.079 | **[0.032, 0.125]** |
| *S. gouazoubira* | effort | 0.425 | **[0.242, 0.704]** |
| *S. gouazoubira* | lunar phase | -0.033 | [-0.076, 0.011] |
| *O. bezoarticus* | Intercept | 0.540 | **[0.382, 0.703]** |
| *O. bezoarticus* | days since maint. | 0.262 | **[0.084, 0.447]** |
| *O. bezoarticus* | Julian day | -0.224 | **[-0.403, -0.050]** |
| *O. bezoarticus* | effort | 0.387 | **[0.169, 0.744]** |
| *O. bezoarticus* | lunar phase | -0.108 | [-0.278, 0.058] |

#### Table S6. Spatiotemporal-avoidance models

**Table S6.** Linear mixed-effects models of log inter-event interval for the Niedballa spatiotemporal-avoidance analysis, for both dyads and across model specifications (baseline; diel-corrected, adding period of day; diel-overlap, adding camera-level overlap; moon-covariate, adding lunar phase). The coefficient of interest is direction (AB vs BA); a coefficient differing from zero indicates directional asymmetry in inter-event intervals. Only the *A. axis / O. bezoarticus* dyad shows a significant direction effect; it remains negative but attenuates under additional covariates, and the adjusted confidence intervals include zero. Station was included as a random intercept. Estimates are on the log scale.

| **Dyad** | **Model** | **Direction (AB) β** | **95% CI** | **p** |
|---|---|---|---|---|
| *Aa* / *Sg* | baseline | 0.085 | [-0.297, 0.467] | 0.662 |
| *Aa* / *Sg* | diel-corrected | 0.078 | [-0.306, 0.463] | 0.689 |
| *Aa* / *Sg* | diel-overlap | 0.077 | [-0.307, 0.462] | 0.693 |
| *Aa* / *Sg* | moon-covariate | 0.072 | [-0.313, 0.457] | 0.714 |
| *Aa* / *Ob* | baseline | -0.445 | [-0.666, -0.223] | **< 0.001** |
| *Aa* / *Ob* | diel-corrected | -0.278 | [-0.643, 0.088] | 0.136 |
| *Aa* / *Ob* | diel-overlap | -0.292 | [-0.658, 0.074] | 0.117 |
| *Aa / Sg* | count (glmmTMB) | 0.005 | [-2.212, 2.223] | 0.996 |
| *Aa / Ob* | count (glmmTMB) | -0.018 | [-0.233, 0.197] | 0.869 |

*Note.* A sensitivity analysis excluding detections within 7 days of a camera-maintenance event confirmed the permutation asymmetry (*A. axis / O. bezoarticus* permutation p < 0.001; *A. axis / S. gouazoubira* contexts p = 0.400–0.609), indicating the asymmetry is not an artefact of human disturbance.

The last two rows (count, glmmTMB) report negative-binomial generalized linear mixed models on the number of directional transitions per station (offset by log effort), on the log-rate scale rather than the log-interval scale; they served as a complementary check and show no directional difference in transition counts for either dyad.

#### Table S7. Return-time models

**Table S7.** Linear mixed-effects models of log return time within detection triples, for both dyads. The coefficient return_sp(Aa) contrasts the return time of *A. axis* with that of the other species; none of the models shows a significant difference, indicating that once an intrusion has occurred, the two species return on comparable timescales. Station was included as a random intercept; log effort and (where indicated) period of day and log intrusion time were included as covariates. Estimates are on the log scale.

| Dyad | Model | return_sp(Aa) β | 95% CI | p |
|---|---|---|---|---|
| *Aa* / *Sg* | baseline | 0.217 | [-0.094, 0.528] | 0.171 |
| *Aa* / *Sg* | diel-corrected | -0.088 | [-0.776, 0.600] | 0.801 |
| *Aa* / *Sg* | intrusion-corrected | 0.111 | [-0.224, 0.445] | 0.517 |
| *Aa* / *Ob* | baseline | 0.304 | [-0.095, 0.704] | 0.135 |
| *Aa* / *Ob* | diel-intrusion | -0.312 | [-0.898, 0.273] | 0.293 |

#### Table S8. Two-species co-occurrence: model selection and the discarded categorical model

**Table S8a.** Model selection (AIC) for the two-species co-occurrence models (occuMulti; Rota et al. 2016) for the *A. axis / S. gouazoubira* dyad. Although the model including the categorical habitat factor (local_hab) had the lowest AIC, it suffered from complete separation (occupancy probabilities driven to 0/1 in sparsely sampled habitat classes), producing non-identifiable parameters with implausibly large standard errors (Table S8b). We therefore report the hab_continuous model – which uses continuous structural covariates and is well-identified – as the primary model in the main text.

| Dyad | Model | AIC | ΔAIC | Status |
|---|---|---|---|---|
| *Aa* / *Sg* | local_hab (categorical) | 27,063.8 | 0.0 | discarded (separation) |
| *Aa* / *Sg* | interax_geo | 27,086.0 | 22.2 | — |
| *Aa* / *Sg* | hab_continuous | 27,130.1 | 66.4 | reported (main text) |
| *Aa* / *Sg* | disturbance | 27,143.2 | 79.5 | — |
| *Aa* / *Sg* | null | 27,184.8 | 121.0 | — |

**Table S8b.** Illustration of the non-identifiability of the categorical-habitat co-occurrence model (local_hab). Selected occurrence coefficients (logit scale) show extreme estimates and standard errors – symptomatic of complete separation – including for the species-interaction term. These values are not interpretable and are shown only to justify the exclusion of this model. The interaction term is reliably estimated in the hab_continuous model ($f3 = -3.69$, $SE = 0.88$, $p < 0.0001$).

| Parameter | Estimate | SE |
|---|---|---|
| [*Aa*] Intercept | 22.77 | 89.90 |
| [*Aa*] Habitat: Grassland | -36.54 | 140.21 |
| [*Sg*] Intercept | 20.21 | 89.90 |
| [*Aa*:*Sg*] Intercept (interaction) | -21.42 | 89.90 |

*Note.* For the *A. axis / O. bezoarticus* dyad, the co-occurrence subset comprised only stations where *A. axis* was present by definition, so *A. axis* occupancy was ≈ 1 at all sites and the interaction term was not estimable (SE > 100); no co-occurrence model is reported for that dyad.

**Table S9.** Nocturnal and lunar activity of the three deer species by invasion level. For each stratum we report the total number of independent detections (*n*, all detections) and the percentage occurring at night (% night), with the resulting activity-pattern class following Azevedo et al. (2018). Lunar columns are based on the subset of nocturnal detections occurring with the moon above the horizon (*n*, moonlit nights): the percentage falling in the full-moon window (lunar phase 3π/4–5π/4) and in the new-moon window, and a qualitative lunar tendency (lunarphilic = detections concentrated near full moon). The two *n* columns therefore count different subsets and are not directly comparable.

| Species | Context | n (all detections) | % night | Activity pattern | n (moonlit nights) | % full moon | % new moon |
|---|---|---|---|---|---|---|---|
| *Aa* | Total invasion | 22,598 | 64.4 | cathemeral | 7,990 | 41.0 | 4.3 |
| *Aa* | Low invasion | 239 | 84.1 | mostly nocturnal | 101 | 38.6 | 1.0 |
| *Aa* | High invasion | 944 | 79.4 | mostly nocturnal | 383 | 33.2 | 2.9 |
| *Aa* | Moderate invasion | 804 | 67.8 | cathemeral | 278 | 37.8 | 1.4 |
| *Aa* | High invasion (Aa/Ob) | 1,038 | 76.3 | mostly nocturnal | 254 | 25.2 | 12.6 |
| *Sg* | No invasion | 6,800 | 61.1 | cathemeral | 2,874 | 64.5 | 1.9 |
| *Sg* | Low invasion | 1,499 | 68.8 | cathemeral | 599 | 39.4 | 1.2 |
| *Sg* | High invasion | 223 | 35.4 | cathemeral | 42 | 7.1 | 0.0 |
| *Sg* | Moderate invasion | 1,100 | 65.2 | cathemeral | 497 | 60.8 | 1.4 |
| *Ob* | High invasion (Aa/Ob) | 840 | 6.0 | diurnal | 30 | 73.3 | 0.0 |

**Table S10.** Return-time dynamics within detection triples by dyad and context. For each triple type we report the number of sequences (n) and the median return time (minutes) – the time taken by the focal species to return after the other species was detected. Note the strongly unequal number of sequences in the *A. axis / O. bezoarticus* dyad (35 vs 107), although the raw numbers of sequences were strongly unequal, the effort-adjusted transition-count model detected no directional difference (Table S6), so the imbalance should be read descriptively rather than as the return-time durations themselves.

| Dyad | Context | Triple (return of) | n | Median return (min) |
|---|---|---|---|---|
| *Aa* / *Ob* | High invasion (Aa/Ob) | *Aa* (Aa→Ob→Aa) | 35 | 1,016 |
| *Aa* / *Ob* | High invasion (Aa/Ob) | *Ob* (Ob→Aa→Ob) | 107 | 905 |
| *Aa* / *Sg* | High invasion | *Aa* (Aa→Sg→Aa) | 102 | 1,046 |
| *Aa* / *Sg* | High invasion | *Sg* (Sg→Aa→Sg) | 25 | 880 |
| *Aa* / *Sg* | Low invasion | *Aa* (Aa→Sg→Aa) | 28 | 1,297 |
| *Aa* / *Sg* | Low invasion | *Sg* (Sg→Aa→Sg) | 85 | 1,336 |
| *Aa* / *Sg* | Moderate invasion | *Aa* (Aa→Sg→Aa) | 78 | 1,323 |
| *Aa* / *Sg* | Moderate invasion | *Sg* (Sg→Aa→Sg) | 82 | 813 |

**Table S11.** Model selection for the hierarchical temporal-partitioning analysis (Iannarilli et al. 2025). For each focal species within each dyad, ΔAIC is the difference in AIC between the model that includes the presence/absence of the co-occurring species and the null model without it; positive values indicate that accounting for the other species improves the fit (i.e., a context-dependent diel shift). Results are shown for both the hierarchical GAM (HGAM, reported in the main text) and the trigonometric formulation (Supplementary). The negative ΔAIC for *O. bezoarticus* indicates that the null model was preferred, providing no support for a detectable activity-curve difference under the fitted model.

| Focal species | Dyad | ΔAIC (HGAM) | ΔAIC (trig.) | Support |
|---|---|---|---|---|
| *Aa* | *Aa* / *Sg* | 525.41 | 469.86 | strong |
| *Sg* | *Aa* / *Sg* | 487.66 | 291.16 | strong |
| *Aa* | *Aa* / *Ob* | 130.81 | 131.01 | strong |
| *Ob* | *Aa* / *Ob* | -17.78 | -6.25 | no supported difference |

**Table S12.** Model selection (WAIC) for single-species occupancy for *A. axis* and *S. gouazoubira*. The six competing occupancy submodels are ranked by WAIC (detection held at the full four-covariate model); ΔWAIC is the difference from the best model. The best-supported model per species (lowest WAIC) is shown in the first row of each block. *O. bezoarticus* is not included: with detections at only four of the seven stations within its endemic range, we did not compare habitat submodels for this species and instead report an intercept-only occupancy estimate (see Methods and main text).

| Species | Occupancy model | WAIC | ΔWAIC |
|---|---|---|---|
| *Aa* | local habitat | 18,394.6 | 0.0 |
| *Aa* | geographic | 18,430.8 | 36.2 |
| *Aa* | landscape 5 km | 18,431.7 | 37.1 |
| *Aa* | disturbance | 18,433.5 | 38.9 |
| *Aa* | null | 18,434.1 | 39.5 |
| *Aa* | landscape 500 m | 18,434.6 | 40.0 |
| *Sg* | local habitat | 10,383.6 | 0.0 |
| *Sg* | disturbance | 10,402.1 | 18.5 |
| *Sg* | geographic | 10,417.2 | 33.6 |
| *Sg* | landscape 500 m | 10,423.5 | 39.9 |
| *Sg* | landscape 5 km | 10,424.5 | 40.9 |
| *Sg* | null | 10,435.7 | 52.2 |

**Table S13.** Occupancy coefficients (posterior median and 95% credible interval, logit scale) from the best-supported single-species models (the local-habitat model) for *A. axis* and *S. gouazoubira*. Coefficients whose interval excludes zero (in bold) indicate a credible effect. Habitat type is relative to the Forest reference level. *O. bezoarticus* was not modeled with habitat covariates (see Table S12 and Methods)

| Species | Covariate | Estimate | 95% CI |
|---|---|---|---|
| *Aa* | Habitat: Forestry | -1.82 | **[-3.32, -0.33]** |
| *Aa* | Habitat: Grassland | -3.91 | **[-5.61, -2.27]** |
| *Aa* | Habitat: Savanna | +1.42 | [-0.71, 3.92] |
| *Aa* | Understory | -1.03 | **[-1.79, -0.25]** |
| *Aa* | Tree cover | -0.38 | [-1.14, 0.36] |
| *Aa* | Rock cover | +0.70 | **[0.08, 1.54]** |
| *Aa* | Water distance | +0.39 | [-0.18, 1.02] |
| *Sg* | Habitat: Forestry | +1.23 | [-0.32, 2.84] |
| *Sg* | Habitat: Grassland | +1.52 | [-0.47, 3.42] |
| *Sg* | Habitat: Savanna | −1.20 | [-3.80, 1.16] |
| *Sg* | Understory | +1.93 | **[0.94, 3.06]** |
| *Sg* | Tree cover | +1.39 | **[0.49, 2.30]** |
| *Sg* | Rock cover | −0.43 | [-1.14, 0.23] |
| *Sg* | Water distance | −0.10 | [-0.77, 0.54] |

**Table S14.** Predicted occupancy (ψ) by habitat type for *A. axis* and *S. gouazoubira*, from the local-habitat models with continuous covariates held at their means. Values illustrate the divergent habitat profiles: *A. axis* peaks in savanna and forest, *S. gouazoubira* in grassland and forestry, with native forest occupied by both. *O. bezoarticus* was not modeled with habitat covariates (see Table S12 and Methods)

| **Habitat** | **ψ *A. axis*** | **ψ *S. gouazoubira*** |
|---|---|---|
| Savanna | 0.95 | 0.14 |
| Forest | 0.83 | 0.33 |
| Forestry | 0.43 | 0.63 |
| Grassland | 0.09 | 0.69 |

**Table S15.** Conditional occupancy (ψ) of each species given the presence or absence of the other, from the primary estimable two-species co-occurrence model for the *A. axis / S. gouazoubira* dyad (occuMulti with continuous habitat covariates). Values are predicted estimates with 95% confidence intervals (bootstrap). The non-overlapping intervals between the 'absent' and 'present' columns are consistent with residual spatial segregation under the fitted model.

| **Focal species** | **Other species absent** | **Other species present** |
|---|---|---|
| *Aa* | 0.80 [0.63, 0.89] | 0.21 [0.09, 0.42] |
| *Sg* | 0.67 [0.51, 0.79] | 0.29 [0.19, 0.47] |

**Table S16.** Annual detections and sampling effort at the eight camera-trap stations of the moderate-invasion level (Durazno), 2019–2025. All eight stations operated continuously from 3 March 2019 to 21 December 2025 without interruption, so camera-nights per year correspond to the number of days the stations were deployed within each year and annual effort is near-constant. RAI is the relative abundance index (independent detections per 100 camera-nights). Habitat covariates at these stations were unchanged over the period. This series is the only direct temporal evidence in the study and underlies the trajectory described in the main text.

| **Year** | **Camera-nights** | ***Aa*** | ***Sg*** | **RAI *Aa*** | **RAI *Sg*** |
|---|---|---|---|---|---|
| 2019 | 2,432 | 90 | 397 | 3.70 | 16.32 |
| 2020 | 2,928 | 77 | 399 | 2.63 | 13.63 |
| 2021 | 2,920 | 92 | 128 | 3.15 | 4.38 |
| 2022 | 2,920 | 126 | 92 | 4.32 | 3.15 |
| 2023 | 2,920 | 127 | 63 | 4.35 | 2.16 |
| 2024 | 2,928 | 145 | 21 | 4.95 | 0.72 |
| 2025 | 2,840 | 147 | 0 | 5.18 | 0 |
| **Total** | **19,880** | **804** | **1,100** | **4.04** | **5.53** |

#### Supplementary Figures

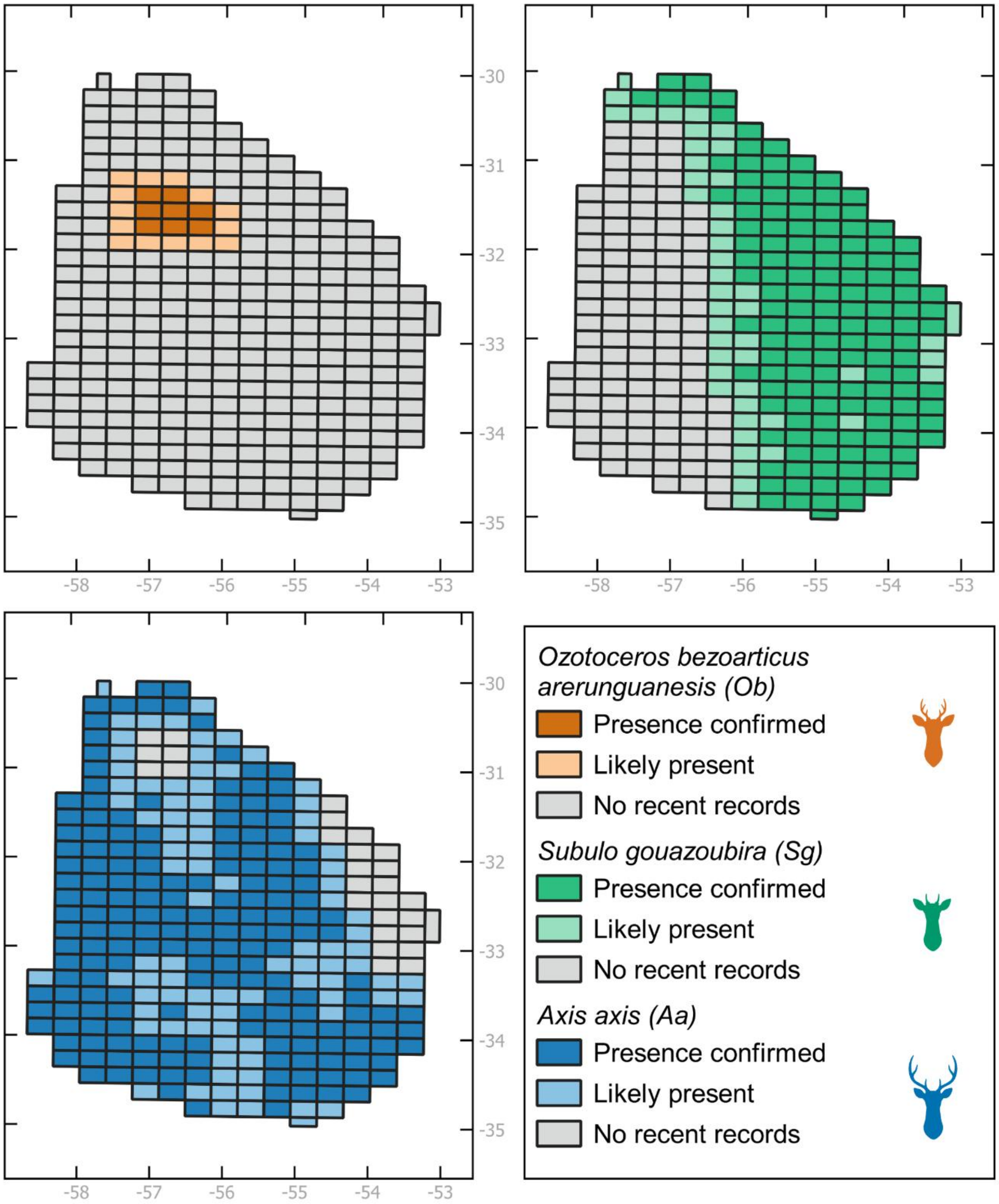


**Figure S1.** Distribution of the three deer species within Uruguay, based on grid cells with confirmed records (from camera-trap data and complementary sources). The map illustrates the broad national distribution of *S. gouazoubira*, the expanding and spatially heterogeneous range of *A. axis* within Uruguay, and the restricted range of *O. bezoarticus arerunguaensis* in the basaltic region of north-central Uruguay (Salto, Paysandú and western Tacuarembó).

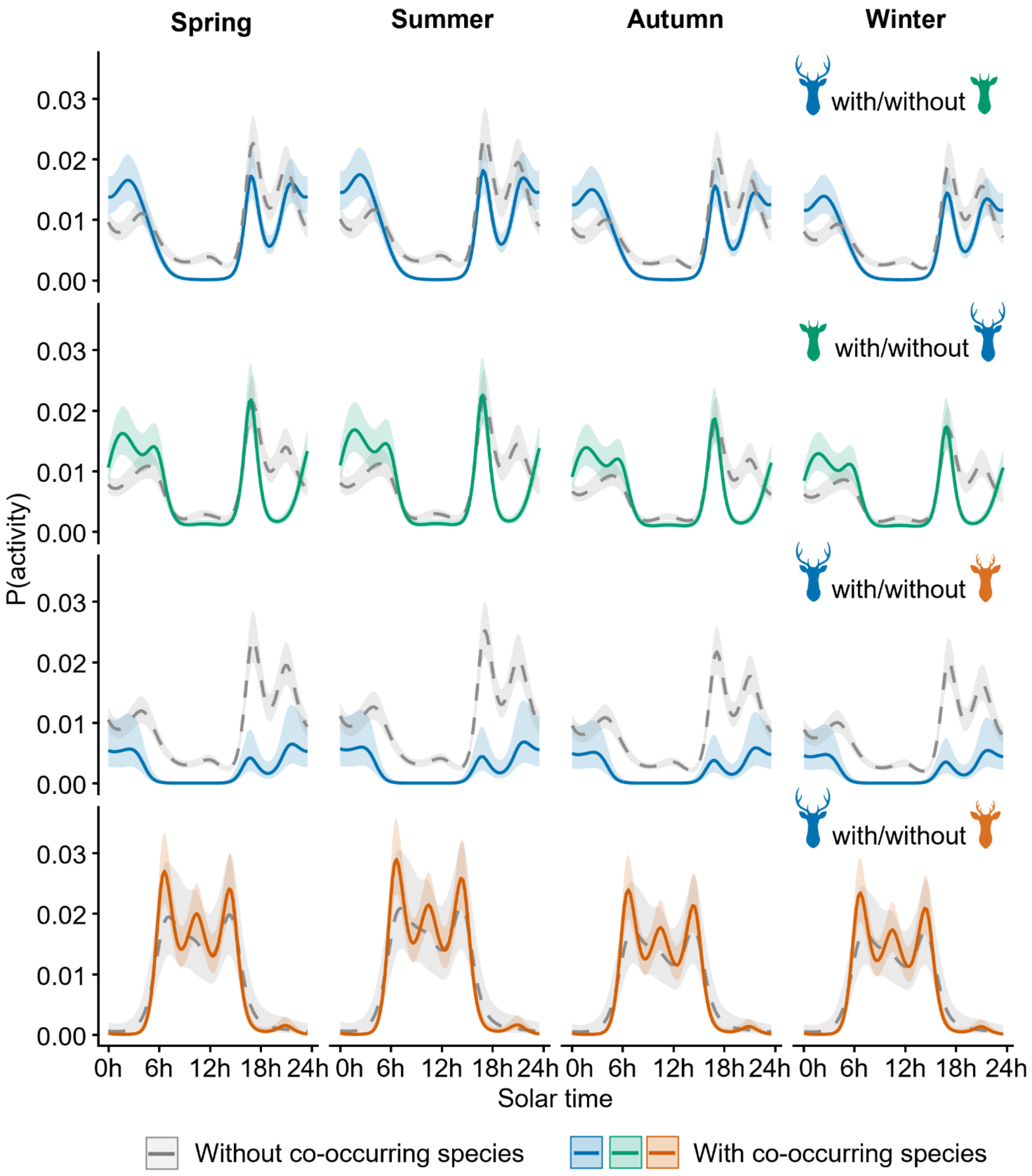


**Figure S2.** Seasonal hierarchical generalized additive models (HGAMs) of diel activity, showing model-predicted activity curves for each species by season (spring, summer, autumn, winter). These complement the season-pooled models in the main text (Figure 6) and indicate whether the diel shifts associated with co-occurrence are consistent across seasons.

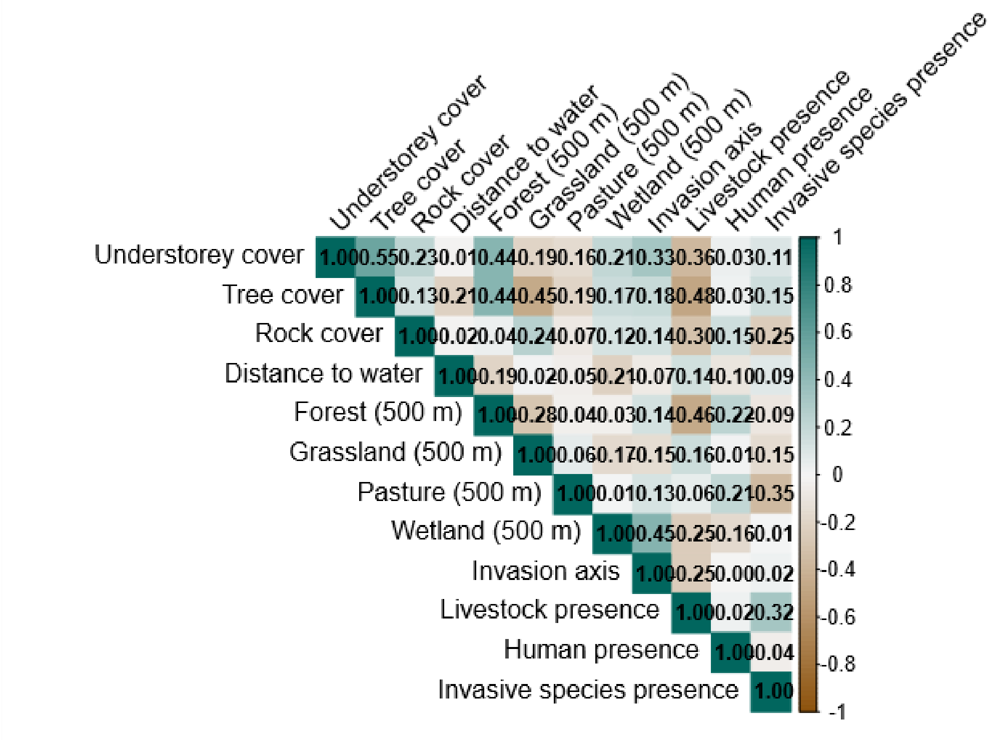


**Figure S3.** Pairwise Pearson correlations among the site-level covariates used in the single-species occupancy models. Cell colour and the printed coefficient give the correlation between each pair of covariates (teal positive, brown negative; the diagonal is 1 by definition). Correlations were inspected to screen for collinearity before model fitting: all pairs fall below |r| = 0.6 except the invasion axis, which is strongly correlated with longitude (r = 0.77) and latitude (r = -0.77) because it summarises the same west-to-east geographic gradient. Longitude and latitude were checked separately and are not shown here; the invasion axis was retained in the models instead, to avoid this redundancy.

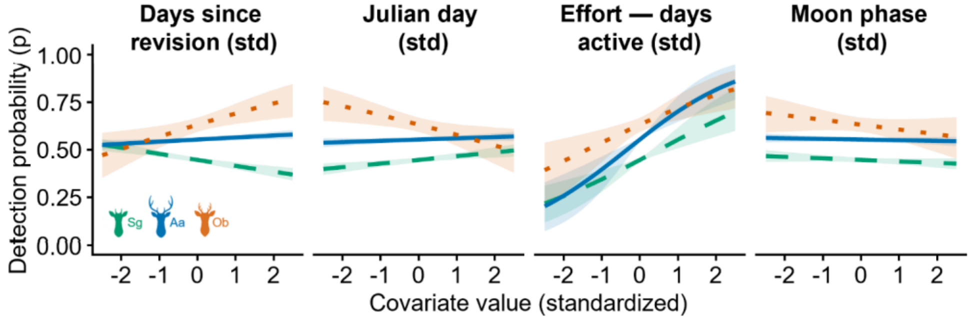


**Figure S4.** Detection probability (p) as a function of the four occasion-level detection covariates – days since the last maintenance or revision, Julian day, within-occasion sampling effort, and mean lunar phase – for the three species, from the full detection submodel retained in each single-species occupancy model (the same detection structure was used in the two-species co-occurrence models). Lines are posterior medians and shaded bands the 95% credible intervals; each covariate is shown on its standardized scale with the remaining covariates held at their mean. Detection increases with sampling effort in all three species; the effect of days since servicing differs in sign among species (see Table S5). Species are identified by colour and line type as in the legend.

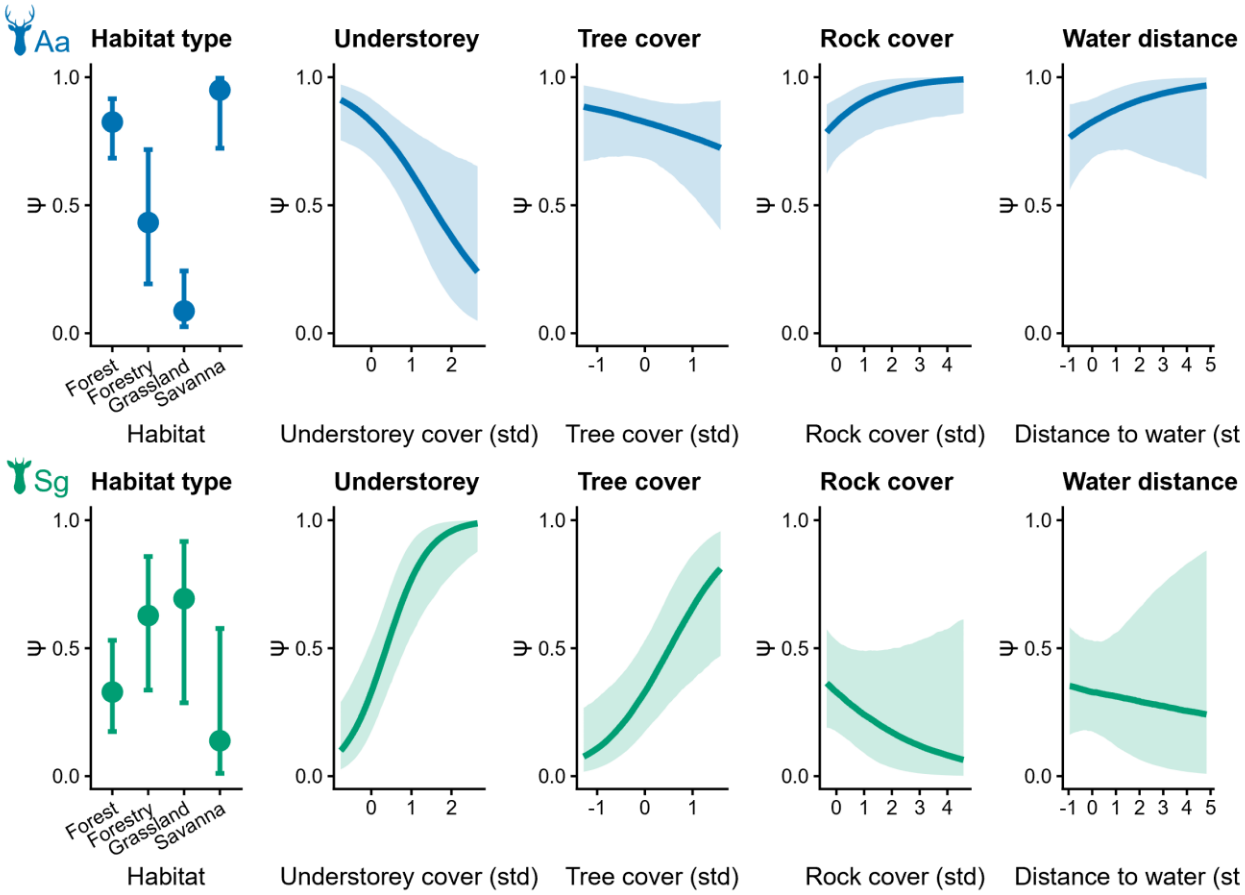


**Figure S5.** Occupancy responses of *A. axis* and *S. gouazoubira* to local-scale covariates, from the best-supported single-species occupancy models (*A. axis* −Aa− and *S. gouazoubira* −Sg−). For categorical habitat, points show predicted occupancy (ψ) by habitat type with 95% credible intervals (other covariates held at their means); for continuous covariates, curves show predicted ψ across the standardized range of each covariate with 95% credible bands. Note the contrasting understory responses of *A. axis* (negative) and *S. gouazoubira* (positive) are the clearest axis of divergence. Species are identified by silhouette and colour as in Figure 1.

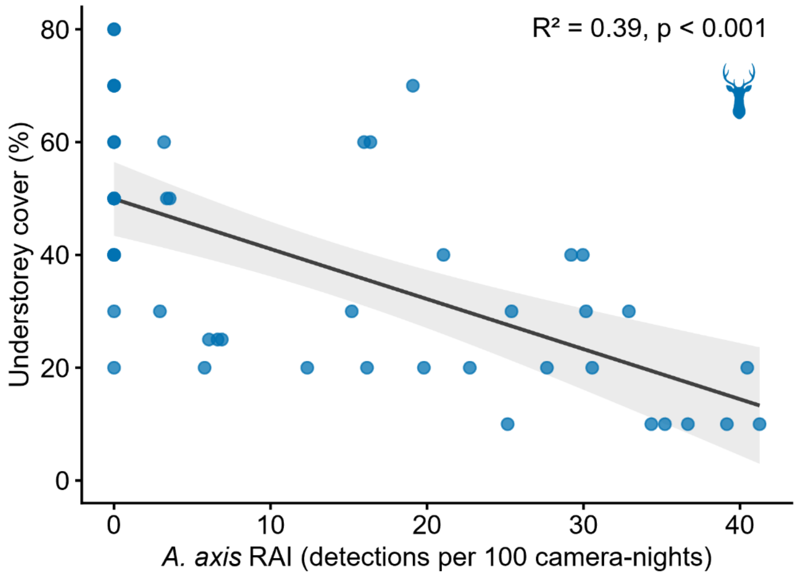


**Figure S6.** Understory cover as a function of *A. axis* relative abundance (RAI) at forest stations only (linear regression β = -0.89, $p < 0.001$, $R^2 = 0.39$; Spearman's ρ = -0.67). Restricting the analysis to forest controls for among-habitat variation in understory. The decline of understory at high *A. axis* abundance is consistent with habitat modification as a possible contributor to spatial segregation, although the cross-sectional design cannot fully separate active degradation from preferential settlement (see Discussion).
